# Local tolerance and innate immune activation of primary human respiratory cells exposed to flagellin

**DOI:** 10.64898/2025.12.19.695341

**Authors:** Guy Barbin, Itziar Sanjuán-García, Mireille Caul-Futy, Christine Caul-Futy, Cindia Ferreira Lopes, Rosy Bonfante, Edouard Sage, Anne-Sophie Lacoste, Jean-Michel Saliou, Delphine Cayet, Ronan MacLoughlin, Mara Baldry, Charlotte Green-Jensen, Arndt G Benecke, Nathalie Heuze Vourc’h, Song Huang, Jean-Claude Sirard, Samuel Constant

**Author notes:** Contributed equally (Jean-Claude Sirard, Samuel Constant). Author contributions statement: Conception and design of the work: J-C.S. and S.C. Acquisition, analysis, and interpretation of the data: I.S-G., M.C-F, R.B., D.C., A-S.L., J-M.S. and A.G.B. Methodology: R.M., C.CF., C.F.L, D.C., and S.H. Resources: E.S. and C.G.J. Supervision: M.B., N.H-V., J-C.S and S.C. Writing – Original draft: G.B., and S.H. Writing – review and editing: G.B., S.H., J-C.S., M.B. Revision and approval of the final version for publication: all authors.

## Abstract

Antibiotic-resistant respiratory infections can lead to treatment failure, highlighting the need for alternative strategies. FLAMOD, a recombinant flagellin, stimulates innate immunity via Toll-like receptor 5 when delivered intranasally. In mice, FLAMOD protects against bacterial pneumonia. The protection is associated with activation of airway epithelial cells. This study aimed to assess the tolerance of human primary respiratory epithelium to FLAMOD administered apically as liquid droplets or by nebulization, to measure the innate immune response, and the pharmacokinetics of FLAMOD. We used epithelia reconstituted from human nasal and bronchial (MucilAir™), small airways (SmallAir™) and alveolar (AlveolAir™) primary epithelial cells, cultured at the air-liquid interface. We report that daily administration of escalating doses of FLAMOD for 5 days was well tolerated by epithelia as barrier integrity, cilia motion and cell viability were not affected. FLAMOD was rapidly degraded without leakage into the basal compartment. Each epithelial model exhibited responses involving pathways of innate defense and immune cell infiltration, which were dose-dependent, with an effective concentration of FLAMOD in the picomolar range. Similar tolerance profile and immune responses were obtained with airway epithelium from cystic fibrosis and chronic obstructive pulmonary disease patients. In conclusion, this study supports the stimulation of epithelial Toll-like receptor 5 signaling to fight against infections of vulnerable patients.

## Introduction

According to the World Health Organization (WHO), acute lower respiratory infections such as pneumonia are the leading cause of death in children and the fourth major cause of death across all ages. Bacterial pneumonia (caused by *Streptococcus pneumoniae* or pneumococcus, *Klebsiella pneumoniae*, *Pseudomonas aeruginosa* and *Haemophilus influenzae*) is responsible for an estimated 4 million deaths annually, with nearly 50% being due to infection with *S. pneumoniae*^1^. The rise of antimicrobial resistance (AMR) has increased the global burden of bacterial pneumonia by complicating treatment and thereby elevating morbidity and mortality. Thus, bacterial infections associated with AMR could become the leading cause of death by 2050^1^, highlighting the urgent need for novel therapeutic strategies. Host-directed therapies, which enhance or modulate the host’s innate immune responses, represent a promising approach^2^.

Toll-like receptors (TLRs) are pattern recognition receptors that detect conserved microbe-derived molecules and consequently activate innate immunity^3^. TLRs are thus targets for host directed therapies and as such, TLR agonists are being investigated as therapeutic agents in infectious diseases^4–6^. Flagellin, the main structural component of bacterial flagellum, is sensed by Toll-like receptor 5 (TLR5) expressed by various cells including epithelial cells (e.g. in the lung or intestine) and myeloid cells^7^. Upon TLR5 binding, flagellin elicits the translocation of NF-κB to the nucleus that, in turn, activates the transcription of genes encoding pro-inflammatory mediators such as C-X-C motif chemokine ligand 8 (CXCL8, also known as interleukin 8, IL-8), C-C motif chemokine ligand 4 (CCL4), CCL5 (also known as RANTES), CCL20, tumor necrosis factor (TNF) and IL-1β^8–11^. In the airways, flagellin elicits the activation and recruitment of immune cells, such as neutrophils, and the production of antimicrobial peptides such as β-defensin^12–15^. The therapeutic potential of flagellin has been evaluated in murine models of pneumonia using intranasal delivery^16–19^. Flagellin prevents pneumonia when administered prior to infection with *P. aeruginosa* or *S. pneumoniae*^13,20^. When administered as an inhaled adjunct to standard-of-care antibiotics amoxicillin or co-trimoxazole, flagellin reduces numbers of pneumococci in the lung and limits dissemination^16,17^. Moreover, flagellin as an adjunct to antibiotic is effective on antibiotic-resistant pneumococcus and reduces the selection of antibiotic-resistant strains^19^. Importantly, antimicrobial activity of gentamicin is enhanced by flagellin against a multidrug-resistant strain of *P. aeruginosa*^21^. Despite the immunostimulation, the lung was not damaged by flagellin adjunct treatment^16^. Finally, nebulized flagellin triggers strong innate immunity in pig airways and boosts antibiotic protection against swine pneumonia caused by *Actinobacillus pleuropneumoniae*^22,23^.

While animal models provide insights into mechanisms and efficacy, translating these findings to humans remains challenging. New Approach Methodologies (NAMs), such as primary differentiated airway epithelial cells, closely mimic respiratory epithelium architecture and innate immune responses, offering high predictive value for respiratory therapeutics. Recent studies showed that flagellin mediates analogous immune-related signaling in human primary epithelium across individuals ^10,11,24–29^. Remarkably, TLR5-mediated activation is also effective when epithelium is infected by *K. pneumoniae, P. aeruginosa* or *S. pneumoniae* ^29–31^. Collectively, these results suggest that human primary epithelium represents a highly potent platform to assess the tolerance and activity to flagellin treatment.

This study evaluated the safety and efficacy of a recombinant flagellin, here named FLAMOD, in human airway epithelial cells cultured at the air-liquid interface (ALI), which closely mimics inhaled therapeutic conditions. FLAMOD was administered apically, either as aerosol via the Aerogen^®^ Solo nebulizer with A-VMN^TM^ mesh (the intended clinical delivery route) or as droplets with a micropipette to mimic intranasal delivery, across nasal, bronchial, small airway, and alveolar models. Tissue integrity, lactate dehydrogenase (LDH) release, and cilia beating activity were monitored as tolerance readouts. FLAMOD showed no toxicity up to 3 µg/cm², was rapidly degraded apically, and did not translocate basally. FLAMOD induced pro-inflammatory gene expression and secretion of immune mediators in all epithelial models.

## Results

### FLAMOD is equally well tolerated by nasal and bronchial epithelia following aerosol and droplet delivery

The effect of exposure to FLAMOD on the respiratory epithelium was evaluated, either by nebulization or liquid droplet deposition at the air interface of nasal models (reconstituted from a pool of 14 healthy donors) and a bronchial model (reconstituted from a single healthy donor). For this purpose, FLAMOD was administered daily for 5 days on the apical surface at doses ranging from 0.0003 µg/cm² to 3 µg/cm² **(Figure 1A**). Control conditions included untreated epithelia and epithelia treated with the vehicle. The first administration of FLAMOD or vehicle started on day 0. Local tolerance was evaluated on days 1, 2, 3, and 4 by analyzing the trans-epithelial electrical resistance (TEER), cilia beating frequency (CBF) and cytotoxicity (LDH release).

**Figure 1.**
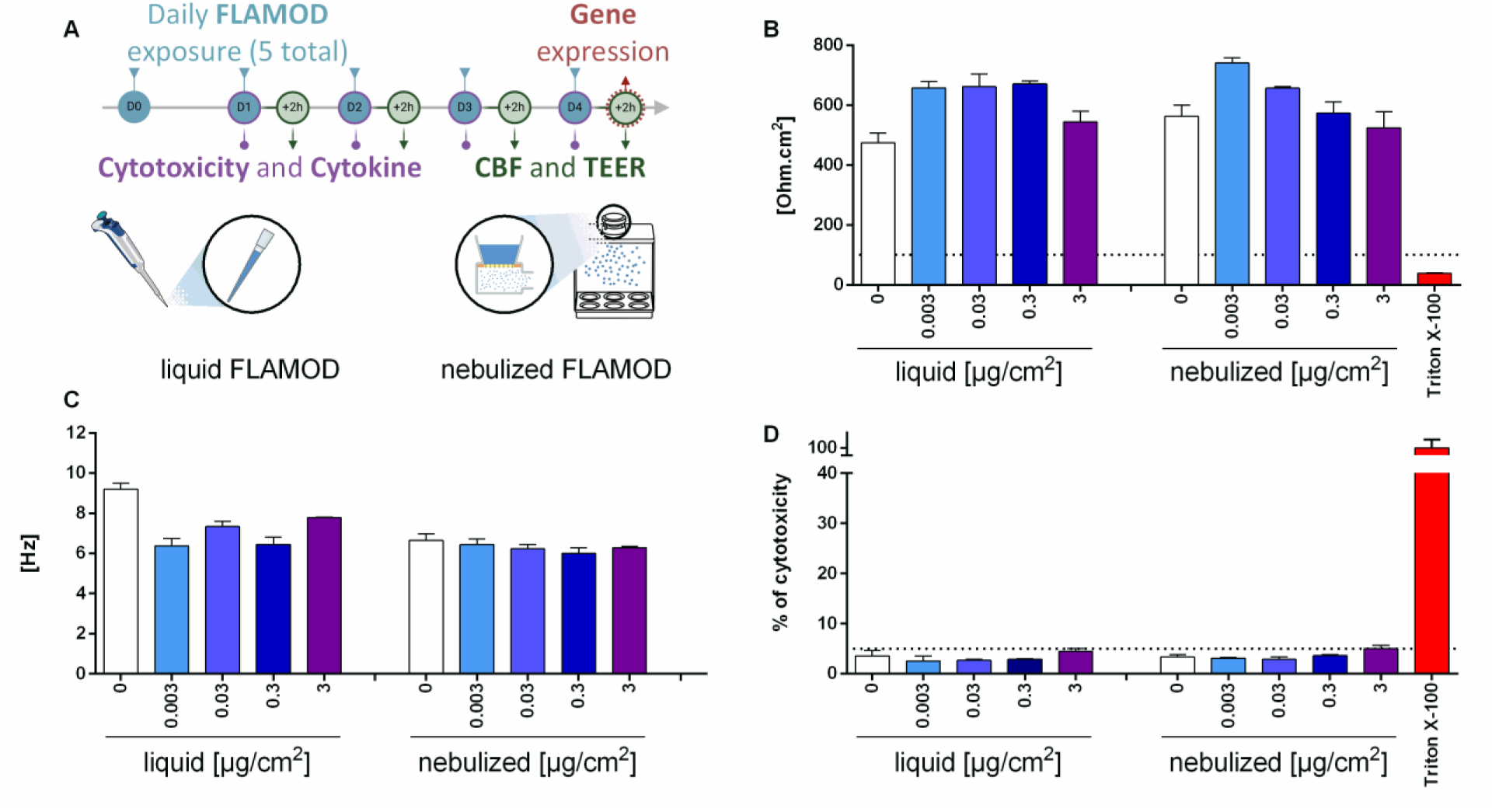
Repeated administration of FLAMOD is well tolerated by nasal epithelium. **(A)** Schematic experimental design. Human nasal epithelial cultures were exposed daily to FLAMOD (administered at specified doses in µg/cm²) or vehicle control, beginning on day 0 and continuing for 4 additional consecutive days (days 1–4). Exposures were applied to the apical surface of the air–liquid interface cultures either via direct pipetting (liquid) or via aerosol delivery (nebulized). Trans-epithelial electrical resistance (TEER), cytotoxicity (LDH release) and cilia beating frequency (CBF) were assessed 2 h post-exposure on each day. On day 4, total RNA was extracted 2 h after the final exposure for gene expression analysis. **(B)** TEER measurements on day 4. TEER value < 100 Ω·cm² indicates compromised epithelial barrier function. (**C)** CBF on day 4. **(D)** Cytotoxicity on day 4. A threshold of 5% LDH release that corresponds to the normal level of cell turnover, was used to define the limit of acceptable cytotoxicity. N=3 replicates; Data are presented as mean ± SEM.

Repeated apical exposure of nasal epithelium to either FLAMOD or vehicle did not affect TEER, which consistently remained above 400 Ω⋅cm^2^ (**Figure 1B; Supplementary Figure 1A**). TEER values were unaffected by the method of delivery, i.e., nebulization or droplet deposition, or the doses of FLAMOD, indicating preserved epithelium tightness. FLAMOD did not trigger any damage on epithelium as LDH values were below the 5% threshold (**Figure 1D**; **Supplementary Figure 2A**), which is associated with the cell turnover. Finaly, cilia function was not altered by FLAMOD as CBF consistently exceeded 4 Hz (**Figure 1D**; **Supplementary Figure 3A**).

In bronchial epithelium, TEER values remained above the 100 Ω**⋅**cm^2^ threshold, indicative of an intact epithelial barrier (**Supplementary Figure 1B**). Although a dose-dependent decrease in TEER was observed, and vehicle-treated samples exhibited unexpectedly low TEER values, no basal-to-apical leakage was detected. Absence of cytotoxicity and preserved ciliary activity further support that FLAMOD does not alter the viability, function and barrier tightness of bronchial epithelium (**Supplementary Figure 2B and 3B**). In conclusion, our findings demonstrate that FLAMOD is well tolerated by both nasal and bronchial epithelial tissues, regardless of the administered dose or the mode of exposure (liquid or nebulized). No evidence of epithelial damage or barrier disruption was observed, supporting the safety profile of FLAMOD for respiratory administration.

### FLAMOD exposure at the air interface is well tolerated by small airways and alveolar epithelium

To investigate responses in the lower respiratory tract, SmallAir™ and AlveolAir™ models were used to mimic the small airways and alveolar epithelium, respectively. As no significant differences in tolerance were observed between FLAMOD delivery methods, exposures were performed exclusively via liquid droplet application at the air-liquid interface, following the protocol described in **Figure 1A**. As observed in nasal and bronchial epithelium, TEER values in SmallAir™ remained stable across repeated FLAMOD exposures (**Supplementary Figure 1C**). In contrast, the alveolar epithelium showed a TEER decrease at the highest FLAMOD doses, suggesting partial tight junction loosening (**Supplementary Figure 1D**). However, no leakage was observed. FLAMOD did not induce any toxicity in the small airways and alveolar epithelium (**Supplementary Figure 2 C-D**). In Small airway, ciliary motion was reduced in a dose-dependent manner (**Supplementary Figure 3C**), but not to ciliopathic levels, as CBF remained above 6 Hz post-exposure, i.e., within the physiological normal range. In summary, our data suggests that repeated FLAMOD administration in small airways and alveolar epithelium is well tolerated.

### FLAMOD stimulates innate immune responses in airway epithelia

As flagellin is associated with TLR5-mediated signaling, the expression of pro-inflammatory genes and the secretion of IL-8 in the basal culture medium was monitored at 2 h and 24 h post-exposure, respectively. Both nebulization and droplet application of FLAMOD induced a comparable, dose-dependent increase in IL-8 secretion in nasal and bronchial epithelia (**Figure 2A; Supplementary Figure 4A-B)**. In contrast, small airway tissue did not secrete IL-8 at any FLAMOD dose (**Supplementary Figure 4C**). Finally, FLAMOD stimulated the secretion of both IL-8 and RANTES in alveolar epithelium (**Supplementary Figure 4D**). In line with the recent observations on gene expression conducted at 4 h post-exposure with liquid droplets of flagellin^29^, our results showed that the transcription of *CCL20, CSF3, DEFB4A* and *IL17C* was upregulated at 2 h post-exposure in nasal and bronchial epithelium following repeated daily FLAMOD stimulation (**Figure 2B-E; Supplementary Figure 5A)**. The activity of FLAMOD was dose-dependent, with an estimated ED50 of 0.005-0.007 µg/cm² (the dose required to elicit half-maximal biological activity; equivalent to 0.2 picomole of FLAMOD/cm²) on both nasal and bronchial epithelium following either liquid or nebulized exposure **(Supplementary Figure 6A-B**). Remarkably, RNA-seq analysis revealed strong similarities in the biological processes induced by FLAMOD regardless the exposure duration (2 h or 4 h), delivery (aerosol or droplet), and tissue type (nasal and bronchial) as shown in **Supplementary Figure 7** and **Supplementary File 1**. A similar expression pattern for *CCL20, CSF3, DEFB4A* and *IL17C* was observed with small airways epithelium exposed to liquid FLAMOD with an estimated ED50 of 0.006 µg/cm² (**Supplementary Figure 5B and Supplementary Figure 6C**). Alveolar tissue also responded to FLAMOD with upregulation of *CCL20* whereas *CSF3, DEFB4A* and *IL17C* were minimally impacted (**Supplementary Figure 5C and Supplementary Figure 6D**). In contrast, transcription of *TNF, TNFAIP6* and *CCL4* was increased. In conclusion, FLAMOD activated innate immune signaling across the whole airway epithelial compartments and demonstrated efficacy at picomolar-range doses regardless of the delivery method. Notably, the epithelial response remained robust and comparable after five repeated doses, indicating sustained responsiveness without signs of desensitization.

**Figure 2.**
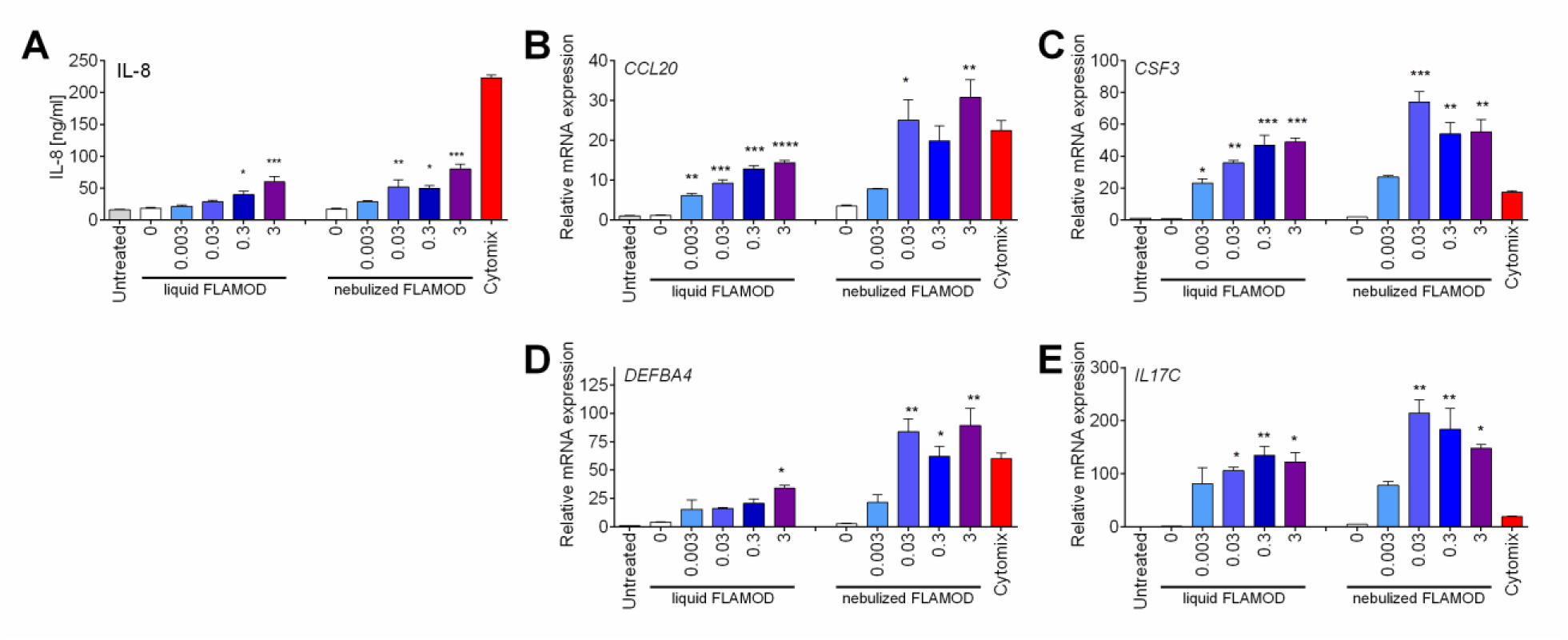
FLAMOD administration by nebulization or liquid droplet stimulates pro-inflammatory responses in human nasal epithelium. Repeated exposure of nasal epithelium to FLAMOD was performed as described in Figure 1 (dose in µg/cm^2^). Cytomix (pro-inflammatory cytokine cocktail) was used as positive control. Tissues were lysed at the end of the experiment for gene expression analysis by RT-qPCR. Secretion of interleukin-8 (IL-8) into the basal compartment was quantified by ELISA on day 4 (24 h accumulation after exposure at day 3). Gene expression was analyzed using RT-qPCR 2 h after the last exposure. **(A)** IL-8 basal secretion on day 4. **(B, C, D and E)** Relative mRNA levels of *CCL20*, *CSF3*, *DEFBA4* and *IL-17C*, respectively. Statistical analysis was performed comparing vehicle control to FLAMOD-treated tissue using one-way ANOVA with Dunnett’s multiple comparison post-test (*p<0.05, **p<0.01, ***p<0.001, ****p<0.0001).

### FLAMOD is rapidly degraded at the air interface of airway epithelium

To further characterize the outcome of FLAMOD in airway epithelium (bronchial MucilAir™), a pharmacokinetics analysis was conducted after single droplet exposure to a high dose of FLAMOD, i.e., 45 µg/cm^2^. At various time points post-exposure, FLAMOD was analyzed and quantified by ELISA and mass spectrometry to assess distribution across compartments: i.e., what was retained in the apical surface (lavages), associated with epithelial cells (cell lysates), and translocated to the basal compartment (basolateral medium). At 30 min post-exposure, FLAMOD levels in apical washes represented less than 10% of the applied dose (**Figure 3A**). By 2 h post-administration, all samples were below the lower limit of quantification (BLLOQ; 3.1 ng/ml) indicating that less than 0.02% of the deposited FLAMOD remained. Similar kinetics were observed in the cell-associated samples with FLAMOD levels being less than 1% of the applied dose at 30 min post-exposure (**Figure 3B**). Importantly, no FLAMOD was detected in the basal culture medium at any time point, indicating no passage through the epithelial barrier (**Figure 3C**). Semi-quantitative LC-MS analyses demonstrated that the detection of FLAMOD-specific peptides aligned with the ELISA data (**Figure 3D-F**). In this context, changes in some proteins in the apical and cell-associated compartment were also observed in response to the FLAMOD treatment (**Supplementary Figure 8**). These findings indicated that FLAMOD undergoes rapid degradation in the apical compartment at the air interface, shows minimal association and internalization by epithelial cells, and does not translocate to the basal compartment. In conclusion, FLAMOD exhibits a short half-life following apical delivery to the airway epithelium but effectively elicits a robust immune response.

**Figure 3.**
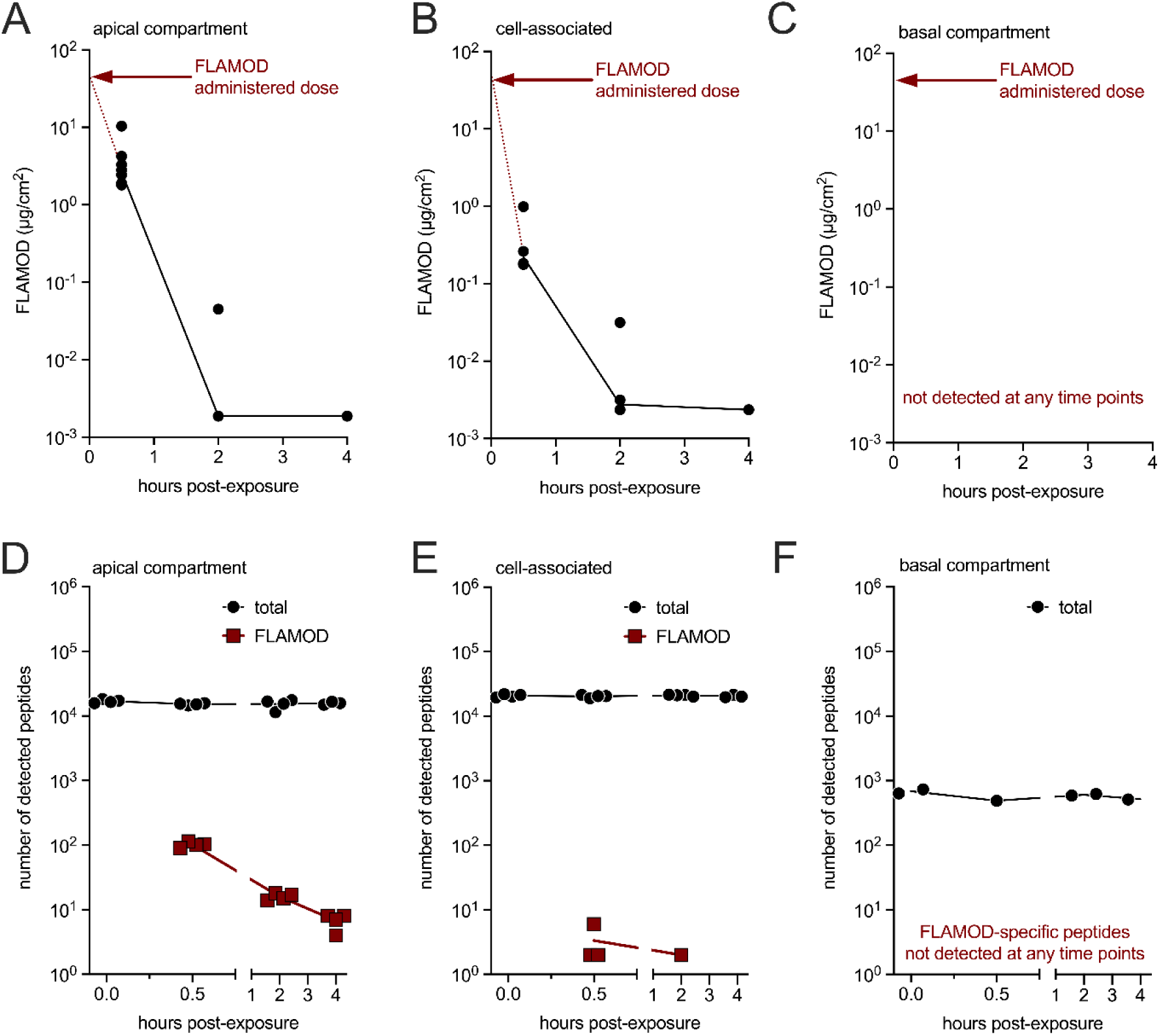
FLAMOD is rapidly degraded following apical delivery to bronchial epithelium. Primary human bronchial epithelium cultured at the air–liquid interface was exposed to high dose of FLAMOD (45 µg/cm^2^) applied directly to the apical surface via pipetting At the indicated time points post-exposure(n=4-8 per time point), samples were collected as follows: apical washes with PBS, lysates of epithelial cells, and basal culture medium to assess the FLAMOD distribution in the apical **(A,D)**, cell-associated **(B, E)** and basal **(C, F)** compartments, respectively. **(A-C)** FLAMOD levels were quantified by ELISA and expressed as mass per surface area. **(D-F)** FLAMOD-specific peptides were analyzed by LC-MS, with total peptide content in samples used for normalization.

### Repeated FLAMOD exposure in airway epithelium from COPD and CF patients shows tolerability and stimulatory profiles comparable to healthy controls

Chronic obstructive pulmonary disease (COPD) and cystic fibrosis (CF) cause structural and functional changes in airway epithelium that weakens immune defenses and increases infections and disease exacerbations^32–35^. The local tolerance and immunomodulatory potential of FLAMOD were therefore evaluated using airway epithelium reconstituted from primary cells isolated from 3 distinct COPD patients and 3 CF patients carrying unrelated mutations of the *CFTR* gene (ΔF508 heterozygous, N1303K heterozygous, and 2184ΔA/W1282X). FLAMOD was administered daily by liquid pipetting (as described in **Figure 1A**) at doses ranging from 0.003-0.3 µg/cm^2^. Similar to airway epithelium from healthy individuals, dose-dependent reduction of TEER was observed in pathological COPD and CF airway epithelium without loss of barrier function (TEER>100 Ω⋅cm^2^ and no leakage of basal culture medium; **Figure 4A**). Repeated administration was well tolerated by both COPD and CF epithelium, with absence of cytotoxicity and conserved cilia function (**Figure 4B-C**). These exposures efficiently stimulated the basal secretion of IL-8 and IL-6 (**Supplementary Figure 9**). FLAMOD-induced effects were less marked in CF epithelium. This more moderate response might be correlated with the different microenvironments in pathological mucus. RNA-seq analysis was performed with epithelium after the fifth exposure to FLAMOD (**Figure 5** and **Supplementary File 2**). When compared to vehicle, exposure to FLAMOD at 0.3 µg/cm^2^ induced the differential expression of 1492, 7834, and 768 genes in healthy, COPD and CF conditions respectively (**Figure 5A**). A comparative analysis as shown in the Venn diagram of **Figure 5B** identified 193 genes common to all conditions. Among the common genes, *CCL20, CSF3, CXCL6, DEFB4A*, *IDO1*, *IL17C, IL23A,* and *TNFAIP6* were upregulated as observed throughout this study (**Figure 5C**). The transcription of the genes *BCL2A1*, *CCL2*, *CHI3L1*, *CXCL5*, *DUOXA2*, *ICAM1*, *IL36A*, *MMP13*, *PGLYRP2*, and *SPRR2F* that also contribute to antimicrobial defenses was also upregulated (**Figure 5D**). Gene ontology analysis on the common 193 genes revealed biological processes induced by FLAMOD that are related to immune response and cytokine signaling (**Figure 5E**). Together, these findings demonstrate that FLAMOD is well tolerated by airway epithelia from COPD and CF patients and exerts a robust immunomodulatory effect by enhancing the expression of genes involved in innate immune defense and antimicrobial responses even in pathological contexts.

**Figure 4.**
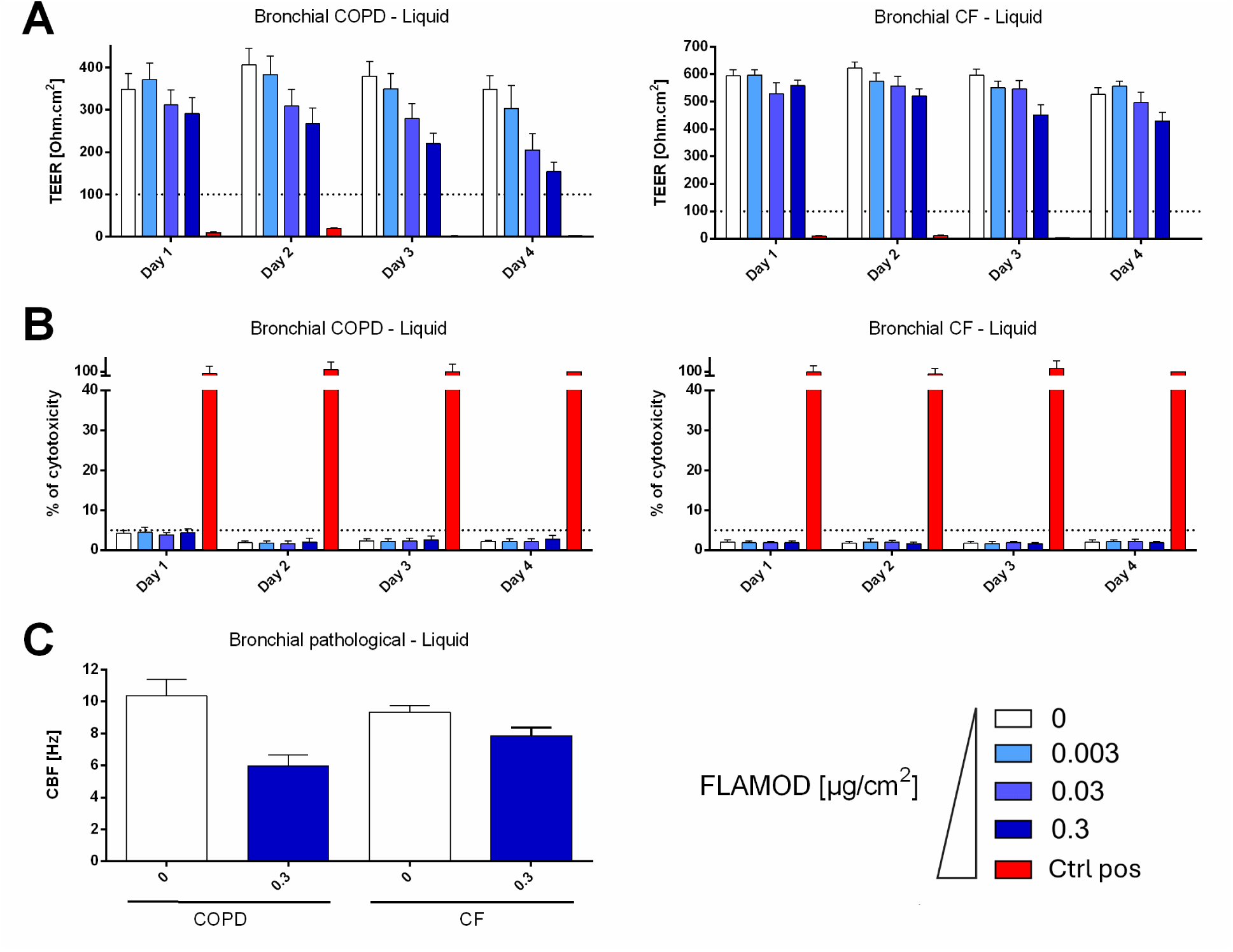
FLAMOD administration is well tolerated by airway epithelium from COPD and CF patients. Bronchial epithelia were reconstituted with primary cells isolated from COPD (left) CF (right) donors and cultured at the air–liquid interface. FLAMOD was administered daily for 5 days by liquid pipetting, as described in Figure 1 (dose in µg/cm^2^). The effects of daily administration on **(A)** tissue integrity (TEER, positive control is Triton X-100), **(B)** cytotoxicity (LDH release positive control is Triton X-100), and **(C)** cilia motion (CBF). TEER measurements < 100 Ω·cm² and cytotoxicity values >5% indicate compromised epithelial barrier function.

**Figure 5.**
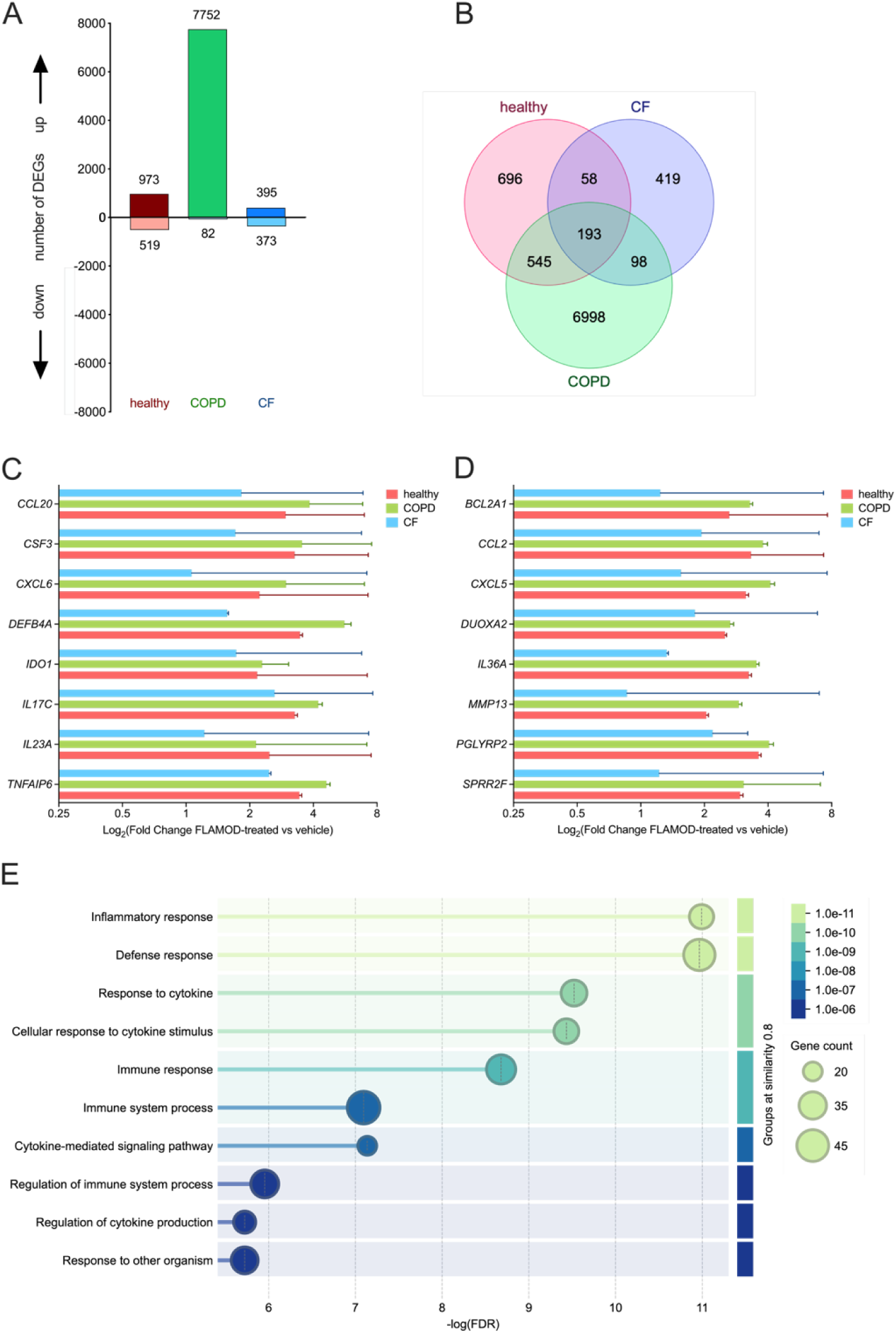
Airway epithelium from COPD and CF patients are stimulated by FLAMOD to promote antimicrobial immune defenses. Bronchial epithelia were reconstituted with primary cells isolated from healthy individuals, COPD, or CF patients and cultured at the air–liquid interface. Vehicle or FLAMOD at 0.3 µg/cm2 was administered daily for 5 days by liquid pipetting, as described in Figure 1. Total RNA was extracted 2 h after the fifth exposure and processed for RNA-seq from experiments. Samples corresponding to repeated dosing of FLAMOD and vehicle buffer were used to analyze the differentially expressed genes (DEG; listed **in Supplementary File 2**). Analysis of the pattern of transcriptional activation was conducted to define the number of genes that were up-regulated and down-regulated by FLAMOD treatment. **(A)** Number of differentially expressed genes (DEG) in various clinical situations. **(B)** Venn diagram illustrating the number of unique and overlapping genes across different conditions. **(C, D)** Fold change of expression of overlapping genes related to innate immune response of healthy airway epithelium and antimicrobial responses. **(E)** Graph of enrichment ontology clusters across overlapping genes.

## Discussion

This study provides first-time evidence that recombinant flagellin FLAMOD is well tolerated by human airway epithelial tissues and elicits a potent innate immune response without compromising epithelial integrity and function. Repeated apical exposures to FLAMOD were evaluated using two delivery methods, aerosol and liquid droplet, with no differences detected between the two approaches. Using physiologically relevant *in vitro* models of the entire human respiratory tract from nasal tissues to alveolar regions, we demonstrated that FLAMOD does not damage the epithelium, as evidenced by the preservation of barrier permeability and ciliary function. Beyond safety, our findings highlighted that the FLAMOD-mediated immunostimulation is dose-dependent, transient and does not trigger desensitization after several exposures. Finally, the safety and stimulatory activity observed in tissues of healthy individuals were also observed in CF and COPD contexts. Altogether, this study supports the development of the use of TLR5 agonist for respiratory delivery with the purpose of triggering local innate defenses and adaptive immunity.

Preservation of TEER values, absence of cytotoxicity, and stable ciliary beating frequency were consistently observed across delivery methods, whether administered as droplets by micropipetting or as aerosolized formulations using a mesh nebulizer. Even at the highest tested concentration (3 µg/cm²), corresponding to approximately 500 times the ED₅₀, FLAMOD did not induce epithelial damage or compromise barrier integrity, highlighting its favorable safety profile in the upper airways^36^. Comparable results were obtained in airway epithelia derived from nasal, bronchial, small airway, and alveolar tissues, further reinforcing the safety of respiratory delivery^37,38^. Supporting its translational relevance, FLAMOD exhibited a similarly benign tolerability profile in epithelial models derived from individuals with CF or COPD, diseases associated with populations susceptible to respiratory infections and in need of interventions that strengthen mucosal defenses without causing inflammation or tissue injury. These advanced airway epithelium models and companion assays adhere to the 3R principles and the FDA Modernization Act 3.0 and enable to deliver a physiologically relevant and ethically sound alternative to animal testing to evaluate the FLAMOD treatment safety (and activity) across the entire respiratory epithelium, as previously described^39^.

Activation of primary bronchial epithelium upon liquid flagellin exposure (isolated from *S. typhimurium* or *P. aeruginosa*) has been previously reported using human primary airway epithelium differentiated from nasal or bronchial cells at air liquid interface ^10,30,31,39,40^, and airway epithelial cell lines^10,24,41,42^. Importantly, the activity stimulated by FLAMOD was analogous when FLAMOD was delivered as aerosol via mesh-nebulizer or as droplets. FLAMOD elicited a robust, dose-dependent activation of innate immune signaling consistent with its TLR5 agonist activity. The estimated ED₅₀ was remarkably low (0.003–0.007 µg/cm², equivalent to approximately 0.2 pmol/cm²), indicating strong potency and supporting its favorable translational potential for clinical application in humans. In nasal, bronchial, and small airway models, FLAMOD induced basal IL-8 secretion and upregulated key pro-inflammatory and antimicrobial genes such as *CCL20, CSF3, DEFB4A*, and *IL17C*, with comparable potency between nebulized and droplet exposures as described recently by Li et al. after a single droplet exposure^29^. Alveolar tissue also responded to FLAMOD, albeit with a distinct transcriptional profile, characterized by increased expression of *CCL20, TNF, TNFAIP6*, and *CCL4*, suggesting cell type-specific differences in TLR5-mediated responses. Importantly, our findings suggest that repeated exposure did not diminish FLAMOD responsiveness, indicating a lack of tolerance or desensitization. Recent studies using epithelial cell lines have shown that flagellin can induce epigenetic reprogramming, thereby altering the cellular response pattern upon subsequent exposure to the same stimulus^42^. In our air–liquid interface airway epithelium model, we did not specifically assess whether five repeated exposures to FLAMOD alter the response compared to a single exposure. However, the overall profile of innate immune and antimicrobial gene expression appeared only moderately affected, suggesting that the observed effects are likely independent of trained immunity mechanisms.

Pharmacokinetic analyses revealed that FLAMOD is rapidly degraded at the air interface of the airway epithelium. Less than 10% of the applied dose remained detectable in the apical compartment 30 min post-administration, and levels fell below the quantification limit by 2 h. Minimal cell-associated FLAMOD and complete absence in the basal medium confirmed rapid local degradation, limited cellular uptake, and no basolateral translocation. These findings suggest that FLAMOD acts transiently and locally, reducing the risk of systemic exposure and off-target effects. The TEER measurements following FLAMOD exposure corroborated the absence of epithelial leakage and suggested minimal systemic exposure. Furthermore, the short half-life of FLAMOD supports the notion that it acts as a “hit-and-run” biologic, that is rapidly recognized by TLR5, promptly initiates downstream signaling, and triggers transient and potent activation of innate immune and antimicrobial defense pathways. This rapid engagement and clearance mechanism minimizes prolonged receptor stimulation, thereby reducing the risk of excessive inflammation while maintaining effective mucosal immune activation.

Activation of pathways related to antimicrobial responses and chemokine signaling has been previously reported following flagellin administration in airway epithelium infected with multi-drug-resistant *K. pneumoniae*^31^. Moreover, the MucilAir™ airway epithelium model has been extensively used to study host–pathogen interactions and evaluate treatments against major bacterial pathogens, including *S. pneumoniae*, *P. aeruginosa, H. influenzae* and *Staphylococcus aureus*, as well as respiratory viruses such as SARS-CoV-2 and influenza viruses^43–46^. Host-directed immunotherapy represents a promising strategy to prevent or treat infections^12^REF). Instead of targeting pathogens, immunotherapy enhances the host’s ability to control infection, either by boosting antimicrobial immune responses or by modulating dysregulated inflammation. Among the key targets are TLRs that detect conserved microbial components and initiate innate immune signaling^3^. By orchestrating both innate and adaptive immune responses, TLRs play a central role in antimicrobial defense. Consequently, Toll-like receptor (TLR) agonists, such as flagellin, have emerged as promising immunotherapeutic agents for the prevention and treatment of infectious diseases^16–23^. When administered to the conducting airways, the airway epithelium serves as the main responder to TLR5-targeted interventions, making it a valuable system for evaluating the therapeutic efficacy of FLAMOD^47–49^. Human primary airway epithelial models, either used alone or reconstituted with immune cells, offer robust platforms to assess the immunostimulatory potential of FLAMOD across diverse pathogenic contexts. These models enable the exploration of FLAMOD both as a standalone therapy and as an adjunct to conventional antimicrobial or antiviral treatment

In conclusion, FLAMOD demonstrates excellent tolerability and efficacy profiles across the entire respiratory epithelium. The ability of FLAMOD to stimulate innate immune defenses locally and transiently, without compromising epithelial integrity, highlights its potential as a mucosal immunomodulator or adjuvant in respiratory therapeutic strategies. Further studies will be essential to confirm these findings and evaluate its clinical efficacy in infectious disease.

## Methods

Detailed protocols, buffer compositions, reagents, media, kits and manufacturers are provided as **Supplementary Materials.**

### FLAMOD preparation

FLAMOD (recombinant flagellin FliC_Δ174-400_ harboring one extra amino acid at the N terminus) is a custom-designed flagellin derived from *Salmonella enterica* serovar Typhimurium FliC^50^. FLAMOD was produced in *Escherichia coli* by the Vaccine Development department (Statens Serum Institut, Denmark), was purified by filtration and chromatography and resuspended in either D-PBS or NaPi buffer (phosphate buffer 10 mM, NaCl 145 mM, polysorbate 80 0.02% (w/v), pH 6.5). Immunostimulatory activity was validated using HEK-Dual™ hTLR5 cells assay. Endotoxin content was assessed with Pyrochrome^®^.

### Exposure of epithelium to FLAMOD

Mucus quantity was normalized by washing epithelia (nasal, bronchial and small airways) 3 days before the first administration of FLAMOD. Solutions of FLAMOD were prepared daily in dedicated buffer (Healthy: D-PBS for nasal/bronchial or NaPi for small airways/alveolar epithelia – COPD and CF: NaPi), which was used as control vehicle. Apical exposures were performed by depositing FLAMOD contained in 10 µl (liquid droplet exposure by pipetting) or by aerosolization (Aerogen^®^ Solo mesh-nebulizer calibrated for deposition equivalent to that of pipetting).

### Trans-Epithelial Electrical Resistance (TEER)

Resistances were measured with a volt-ohm-meter after the addition of 200 μl of NaCl buffer to the apical compartment. Control for loss of barrier function was obtained by apical treatment with 100 µl of Triton X-100 (10% v/v). Resistances were converted to TEER using the following equation: TEER (Ω·cm^2^) = [resistance measured (Ω) – resistance membrane (Ω)]*membrane surface (cm^2^); Where the resistance of the membrane = 100 Ω.

### Lactate dehydrogenase (LDH)

Cytotoxicity was evaluated by LDH release in the basal medium using a dedicated kit. Control for 100% cytotoxicity was obtained by apical treatment with 100 µl of Triton X-100 (10% v/v).

### Cilia Beating Frequency (CBF)

CBF was measured by capturing 256 images at 125 frames per second at 34°C. CBF was then calculated and expressed as Hz using Cilia-X software.

### ELISA

Interleukin-8 and RANTES were measured in the basal medium using dedicated ELISA kits. FLAMOD was measured using custom-made ELISA.

### Gene expression

Cultures were washed with PBS and lysed with RA1 buffer complemented with 2% TCEP. Total RNA was extracted with the Nucleospin RNA kit and reverse-transcribed with the High-Capacity cDNA Archive Kit. The cDNA was amplified using TaqMan or SYBR green-based RTqPCR (**Supplementary Table 3**). Relative mRNA levels were determined using the ΔΔCq method. Transcriptome analyses using RNAseq as described in Supplementary Materials.

### Mass spectrometry analysis

Samples from apical washes, cell lysates, and basal culture medium were analyzed to determine the frequency and composition of peptides derived from FLAMOD and human proteins.

### Statistical analysis

Data were expressed as mean±SEM and assumed to have normal distribution. Differences between two or more groups were tested by two-way ANOVA with Dunnett’s multiple comparison post-tests. The P values <0.05 were considered statistically significant (*p<0.05, **p<0.01, ***p<0.001, ****p<0.0001).

## Acknowledgements

We thank Laure Lemée and George Michel Haustant from Biomics Platform, C2RT, Institut Pasteur, Paris, France for RNA-seq that are supported by France Génomique (ANR-10-INBS-09) and IBISA.

## References

1 Murray CJL, Ikuta KS, Sharara F, Swetschinski L, Aguilar GR, Gray A, et al. Global burden of bacterial antimicrobial resistance in 2019: a systematic analysis. The Lancet 2022;399:629–55. 10.1016/S0140-6736(21)02724-0.

2 Wallis RS, O’Garra A, Sher A, Wack A. Host-directed immunotherapy of viral and bacterial infections: past, present and future. Nat Rev Immunol 2023;23:121–33. 10.1038/s41577-022-00734-z.

3 Fitzgerald KA, Kagan JC. Toll-like Receptors and the control of immunity. Cell 2020;180:1044–66. 10.1016/j.cell.2020.02.041.

4 Duggan JM, You D, Cleaver JO, Larson DT, Garza RJ, Guzmán Pruneda FA, et al. Synergistic interactions of TLR2/6 and TLR9 induce a high level of resistance to lung infection in mice. J Immunol 2011;186:5916–26. 10.4049/jimmunol.1002122.

5 Tuvim MJ, Gilbert BE, Dickey BF, Evans SE. Synergistic TLR2/6 and TLR9 Activation Protects Mice against Lethal Influenza Pneumonia. PLoS One 2012;7:e30596. 10.1371/journal.pone.0030596.

6 Yang J, Yan H. TLR5: beyond the recognition of flagellin. Cell Mol Immunol 2017;14:1017–9. 10.1038/cmi.2017.122.

7 Vijayan A, Rumbo M, Carnoy C, Sirard J-C. Compartmentalized Antimicrobial Defenses in Response to Flagellin. Trends in Microbiology 2017;9:143. 10.1016/j.tim.2017.10.008.

8 Murthy KGK, Deb A, Goonesekera S, Szabó C, Salzman AL. Identification of conserved domains in Salmonella muenchen flagellin that are essential for its ability to activate TLR5 and to induce an inflammatory response in vitro. J Biol Chem 2004;279:5667–75. 10.1074/jbc.M307759200.

9 Gewirtz AT, Navas TA, Lyons S, Godowski PJ, Madara JL. Cutting edge: bacterial flagellin activates basolaterally expressed TLR5 to induce epithelial proinflammatory gene expression. J Immunol 2001;167:1882–5. 10.4049/jimmunol.167.4.1882.

10 Losol P, Ji M-H, Kim JH, Choi J-P, Yun J-E, Seo J-H, et al. Bronchial epithelial cells release inflammatory markers linked to airway inflammation and remodeling in response to TLR5 ligand flagellin. World Allergy Organ J 2023;16:100786. 10.1016/j.waojou.2023.100786.

11 Wang R, Ahmed J, Wang G, Hassan I, Strulovici-Barel Y, Salit J, et al. Airway Epithelial Expression of TLR5 Is Downregulated in Healthy Smokers and Smokers with Chronic Obstructive Pulmonary Disease. The Journal of Immunology 2012;189:2217–25. 10.4049/jimmunol.1101895.

12 Matarazzo L, Costa C, Porte R, Saliou J-M, Figeac M, Delahaye F, et al. Neutrophil subsets enhance the efficacy of host-directed therapy in pneumococcal pneumonia. Mucosal Immunol 2025;18:257–68. 10.1016/j.mucimm.2024.11.009.

13 Muñoz N, Van Maele L, Marqués JM, Rial A, Sirard J-C, Chabalgoity JA. Mucosal administration of flagellin protects mice from Streptococcus pneumoniae lung infection. Infect Immun 2010;78:4226–33. 10.1128/IAI.00224-10.

14 Janot L, Sirard J-C, Secher T, Noulin N, Fick L, Akira S, et al. Radioresistant cells expressing TLR5 control the respiratory epithelium’s innate immune responses to flagellin. European Journal of Immunology 2009;39:1587–96. 10.1002/eji.200838907.

15 Honko AN, Sriranganathan N, Lees CJ, Mizel SB. Flagellin is an effective adjuvant for immunization against lethal respiratory challenge with Yersinia pestis. Infect Immun 2006;74:1113–20. 10.1128/IAI.74.2.1113-1120.2006.

16 Porte R, Fougeron D, Muñoz Wolf N, Tabareau J, Georgel A-F, Wallet F, et al. A Toll-Like Receptor 5 Agonist Improves the Efficacy of Antibiotics in Treatment of Primary and Influenza Virus-Associated Pneumococcal Mouse Infections. Antimicrobial Agents and Chemotherapy 2015. 10.1128/AAC.01210-15.

17 Matarazzo L, Casilag F, Porte R, Wallet F, Cayet D, Faveeuw C, et al. Therapeutic Synergy Between Antibiotics and Pulmonary Toll-Like Receptor 5 Stimulation in Antibiotic-Sensitive or -Resistant Pneumonia. Front Immunol 2019;10:723. 10.3389/fimmu.2019.00723.

18 Baldry M, Costa C, Zeroual Y, Cayet D, Pardessus J, Soulard D, et al. Targeted delivery of flagellin by nebulization offers optimized respiratory immunity and defense against pneumococcal pneumonia. Antimicrob Agents Chemother 2024;68:e0086624. 10.1128/aac.00866-24.

19 Costa C, Sirard J-C, Gibson PS, Veening J-W, Gjini E, Baldry M. Triggering Toll-Like Receptor 5 Signaling During Pneumococcal Superinfection Prevents the Selection of Antibiotic Resistance. J Infect Dis 2024;230:e1126–35. 10.1093/infdis/jiae239.

20 Yu F, Cornicelli MD, Kovach MA, Newstead MW, Zeng X, Kumar A, et al. Flagellin Stimulates Protective Lung Mucosal Immunity: Role of Cathelicidin Related Antimicrobial Peptide (CRAMP). J Immunol 2010;185:1142–9. 10.4049/jimmunol.1000509.

21 Maia AR, Cezard A, Fouquenet D, Vasseur V, Briard B, Sirard J-C, et al. Preventive nasal administration of flagellin restores antimicrobial effect of gentamicin and protects against a multidrug-resistant strain of Pseudomonas aeruginosa. Antimicrobial Agents and Chemotherapy 2024;68:e01361–23. 10.1128/aac.01361-23.

22 Baldry M, Caballero I, Ruuls L, Cayet D, Zeroual Y, Costa C, et al. Flagellin nebulization enhances respiratory immune responses in the porcine model 2025:2025.03.17.643810. 10.1101/2025.03.17.643810.

23 Fleurot I, Guillory V, Barc C, Riou M, Deslis A, Pléau A, et al. Flagellin aerosol administration improves the efficacy of antibiotic treatment in Actinobacillus pleuropneumoniae infected pigs 2025:2025.03.17.643685. 10.1101/2025.03.17.643685.

24 Adamo R, Sokol S, Soong G, Gomez MI, Prince A. Pseudomonas aeruginosa Flagella Activate Airway Epithelial Cells through asialoGM1 and Toll-Like Receptor 2 as well as Toll-Like Receptor 5. Am J Respir Cell Mol Biol 2004;30:627–34. 10.1165/rcmb.2003-0260OC.

25 Bigot J, Ruffin M, Guitard J, Vellaissamy S, Thorez S, Corvol H, et al. Effect of Flagellin Pre-Exposure on the Inflammatory and Antifungal Response of Bronchial Epithelial Cells to Fungal Pathogens. J Fungi (Basel) 2022;8:1268. 10.3390/jof8121268.

26 Bigot J, Guillot L, Guitard J, Ruffin M, Corvol H, Chignard M, et al. Respiratory Epithelial Cells Can Remember Infection: A Proof-of-Concept Study. J Infect Dis 2020;221:1000–5. 10.1093/infdis/jiz569.

27 Ramirez-Moral I, Ferreira BL, Butler JM, van Weeghel M, Otto NA, de Vos AF, et al. HIF-1α Stabilization in Flagellin-Stimulated Human Bronchial Cells Impairs Barrier Function. Cells 2022;11:391. 10.3390/cells11030391.

28 Arnason JW, Murphy JC, Kooi C, Wiehler S, Traves SL, Shelfoon C, et al. Human β-defensin-2 production upon viral and bacterial co-infection is attenuated in COPD. PLOS ONE 2017;12:e0175963. 10.1371/journal.pone.0175963.

29 Li X, Cayet D, Zeroual Y, Sanjuán-García I, Bonnefond A, Derhourhi M, et al. Flagellin-mediated TLR5 activation enhances innate immune responses in healthy and diseased human airway epithelium 2025:2025.11.06.686934. 10.1101/2025.11.06.686934.

30 Ramirez-Moral I, Yu X, Butler JM, Weeghel M van, Otto NA, Ferreira BL, et al. mTOR-driven glycolysis governs induction of innate immune responses by bronchial epithelial cells exposed to the bacterial component flagellin. Mucosal Immunology 2021;14:594–604. 10.1038/s41385-021-00377-8.

31 van Linge CCA, Hulme KD, Peters-Sengers H, Sirard J-C, Goessens WHF, de Jong MD, et al. Immunostimulatory Effect of Flagellin on MDR-Klebsiella-Infected Human Airway Epithelial Cells. International Journal of Molecular Sciences 2024;25:309. 10.3390/ijms25010309.

32 De Rose V, Molloy K, Gohy S, Pilette C, Greene CM. Airway Epithelium Dysfunction in Cystic Fibrosis and COPD. Mediators of Inflammation 2018;2018:1309746. 10.1155/2018/1309746.

33 Gao W, Li L, Wang Y, Zhang S, Adcock IM, Barnes PJ, et al. Bronchial epithelial cells: The key effector cells in the pathogenesis of chronic obstructive pulmonary disease? Respirology 2015;20:722–9. 10.1111/resp.12542.

34 Fenker DE, McDaniel CT, Panmanee W, Panos RJ, Sorscher EJ, Sabusap C, et al. A Comparison between Two Pathophysiologically Different yet Microbiologically Similar Lung Diseases: Cystic Fibrosis and Chronic Obstructive Pulmonary Disease. International Journal of Respiratory and Pulmonary Medicine 2018;5:098. 10.23937/2378-3516/1410098.

35 Sethi S. Infection as a comorbidity of COPD. Eur Respir J 2010;35:1209–15. 10.1183/09031936.00081409.

36 de Ávila RI, Müller I, Barlow H, Middleton AM, Theiventhran M, Basili D, et al. Evaluation of a non-animal toolbox informed by adverse outcome pathways for human inhalation safety. Front Toxicol 2025;7:1426132. 10.3389/ftox.2025.1426132.

37 Huang S, Boda B, Vernaz J, Ferreira E, Wiszniewski L, Constant S. Establishment and characterization of an *in vitro* human small airway model (SmallAir^TM^). European Journal of Pharmaceutics and Biopharmaceutics 2017;118:68–72. 10.1016/j.ejpb.2016.12.006.

38 Lopes CF, Laurent E, Caul-Futy M, Dubois J, Mialon C, Chojnacki C, et al. A Novel In Vitro Primary Human Alveolar Model (AlveolAir^TM^) for H1N1 and SARS-CoV-2 Infection and Antiviral Screening. Microorganisms 2025;13:572. 10.3390/microorganisms13030572.

39 Ramirez-Moral I, Schuurman AR, van Linge CCA, Butler JM, Yu X, de Haan K, et al. Single-cell transcriptomics reveals subset-specific metabolic profiles underpinning the bronchial epithelial response to flagellin. iScience 2024;27:110662. 10.1016/j.isci.2024.110662.

40 Li P, Sheng Q, Huang L-C, Turner JH. Epithelial innate immune response to Pseudomonas aeruginosa-derived flagellin in chronic rhinosinusitis. Int Forum Allergy Rhinol 2023;13:1937– 48. 10.1002/alr.23164.

41 Blohmke CJ, Victor RE, Hirschfeld AF, Elias IM, Hancock DG, Lane CR, et al. Innate immunity mediated by TLR5 as a novel antiinflammatory target for cystic fibrosis lung disease. J Immunol 2008;180:7764–73. 10.4049/jimmunol.180.11.7764.

42 Bigot J, Legendre R, Hamroune J, Jacques S, Le Gars M, Millet N, et al. The epigenomic landscape of bronchial epithelial cells reveals the establishment of trained immunity. Genes Immun 2025:1–12. 10.1038/s41435-025-00357-z.

43 Maisonnasse P, Guedj J, Contreras V, Behillil S, Solas C, Marlin R, et al. Hydroxychloroquine use against SARS-CoV-2 infection in non-human primates. Nature 2020;585:584–7. 10.1038/s41586-020-2558-4.

44 Jonsdottir HR, Siegrist D, Julien T, Padey B, Bouveret M, Terrier O, et al. Molnupiravir combined with different repurposed drugs further inhibits SARS-CoV-2 infection in human nasal epithelium in vitro. Biomed Pharmacother 2022;150:113058. 10.1016/j.biopha.2022.113058.

45 Do TND, Abdelnabi R, Boda B, Constant S, Neyts J, Jochmans D. The triple combination of Remdesivir (GS-441524), Molnupiravir and Ribavirin is highly efficient in inhibiting coronavirus replication in human nasal airway epithelial cell cultures and in a hamster infection model. Antiviral Research 2024;231:105994. 10.1016/j.antiviral.2024.105994.

46 Boda B, Benaoudia S, Huang S, Bonfante R, Wiszniewski L, Tseligka ED, et al. Antiviral drug screening by assessing epithelial functions and innate immune responses in human 3D airway epithelium model. Antiviral Res 2018;156:72–9. 10.1016/j.antiviral.2018.06.007.

47 Van Maele L, Fougeron D, Janot L, Didierlaurent A, Cayet D, Tabareau J, et al. Airway structural cells regulate TLR5-mediated mucosal adjuvant activity. Mucosal Immunol 2014;7:489–500. 10.1038/mi.2013.66.

48 Fleurot I, López-Gálvez R, Barbry P, Guillon A, Si-Tahar M, Bähr A, et al. TLR5 signalling is hyper-responsive in porcine cystic fibrosis airways epithelium. J Cyst Fibros 2022;21:e117–21. 10.1016/j.jcf.2021.08.002.

49 López-Gálvez R, Fleurot I, Chamero P, Trapp S, Olivier M, Chevaleyre C, et al. Airway Administration of Flagellin Regulates the Inflammatory Response to Pseudomonas aeruginosa. Am J Respir Cell Mol Biol 2021;65:378–89. 10.1165/rcmb.2021-0125OC.

50 Nempont C, Cayet D, Rumbo M, Bompard C, Villeret V, Sirard J-C. Deletion of flagellin’s hypervariable region abrogates antibody-mediated neutralization and systemic activation of TLR5-dependent immunity. J Immunol 2008;181:2036–43. 10.4049/jimmunol.181.3.2036.

