## Supplementary materials for "Local tolerance and innate immune activation of primary human respiratory cells exposed to flagellin"

[*] Contributed equally

The supplementary materials include:

- Supplementary methods
- 9 supplementary figures
- 3 supplementary tables
- 2 supplementary files (as separate files that list the differentially expressed genes)

### Supplementary methods

The list of chemicals, media, antibodies, kits, equipment, software and deposited data are provided in **Supplementary Table 1**.

*3D ALI cultures (MucilAir™, SmallAir™ and AlveolAir™)*

All models are commercially available and were provided by Epithelix Sàrl. MucilAir™ (nasal and bronchial) and SmallAir™ (small airways) are pseudostratified and ready-to-use 3D models of human airway epithelium, constituted with primary human epithelial cells freshly isolated from surgical pieces. When switched at the air-liquid interface, the progenitor cells undergo a progressive differentiation and polarization to an electrically tight, mucus producing, fully ciliated epithelia. The mature MucilAir™ is composed of basal cells, ciliated cells and goblet cells. SmallAir™ exhibits similar features and cellular composition, with the additional presence of club cells. AlveolAir™ is composed of type I and type II pneumocytes co-cultured with primary human lung endothelial cells. Epithelia were cultured in their dedicated medium and kept at 37°C, 5% $CO_2$, 100% humidity. Nasal epithelia were reconstituted from a pool of 14 different donors. Bronchial, small airways and alveolar models were reconstituted from unique donors. Pathological epithelia were reconstituted from patients with cystic fibrosis (CF, donor 1: ΔF508 hetero, donor 2: N1303K hetero, donor 3: 2184ΔA/W1282X) or chronic obstructive pulmonary disease (COPD). Details for each culture is provided in **Supplementary Table 2**.

*Buffers and solutions*

| | |
|---|---|
| NaPi Buffer | 10 mM phosphate, 145 mM NaCl, polysorbate 80 0.02 % (w/v) pH 6.5 |
| NaCl Buffer | NaCl 0.9 %, HEPES 10 mM, $CaCl_2$ 1.25 mM |
| Triton-X100 | 10% (v/v) in NaCl 0.9% |
| Cytomix | 500 ng/ml TNF, 200 µg/ml LPS, FCS 1% (v/v) |

*Administration of FLAMOD by aerosolization*

Exposures were performed with a heated exposure chamber (Vitrocell® Cloud alpha12) with a mesh nebulizer (Aerogen® Solo nebulizer with A-VMN™ mesh). The system was completely washed, primed with 400 µl of PBS and calibrated using the integrated microbalance to reach desired mass deposition/cm$^2$. Wells in the exposure chambers were loaded with MucilAir™ culture medium and heated at 37°C. Four hundred microliters of FLAMOD or vehicle were then loaded into the mesh nebulizer and cultures were placed in the exposure chamber. Complete nebulization was performed, generating a cloud that was let for 5 minutes to ensure complete deposition onto the apical surface. Cultures were then transferred back into the incubators (37°C, 5% $CO_2$, 100% humidity).

*FLAMOD-specific ELISA*

Maxisorp 96 well plates (Nunc) were coated with 50 µl of capture anti-FLAMOD monoclonal antibody (Mab, 9H10-R2-2E8 mouse IgG1 - lot vt220802-7034 - Biotem) at 5 µg/ml in PBS (D-PBS w/o Ca/Mg, GIBCO) and incubated overnight at 4°C. The plates were washed 3 times with PBS supplemented with Tween 20 (ITW Reagent, 0.05% Tween 20, PBS-T) and saturated for 2 h at room temperature (RT) with 300 µl per well ELISA/ELISPOT diluent 1X (Invitrogen, 5X solution diluted in sterile water). The plates were washed again 3 times with PBS/T. Samples were diluted 1:5 in ELISA/ELISPOT diluent and were distributed per well in the microplates. To generate a calibration curve, FLAMOD (lot 4182.5) was diluted from 100 ng/ml to 0.78 ng/ in ELISA/ELISPOT diluent and 50 µl were distributed per well. Plates were incubated for 2 h at RT and washed 3 times with PBS-T. Fifty microliters per well of the biotinylated FLAMOD-specific detection Mab (4C1H7 mouse IgG1 - lot b220809-7034 - Biotem) diluted at 2 µg/ml in ELISA/ELISPOT diluent were distributed and incubated for 1 h at RT. Plates were washed 3 times with PBS-T and 50 µl per well of the avidin-HRP diluted 1:2000 in ELISA/ELISPOT diluent were added and incubated for 25 min at RT. Plates were washed 5 times with PBS-T. The reaction was developed using 50 µl per well of the TMB substrate reagent (Interchim). Plates were incubated for 5 min at RT in the dark, prior to stopping the reaction by adding 25 µl $H_2SO_4$ 2N (Sigma) per well. The plates were read immediately with a Multiskan FC (Thermo Fisher Scientific) at 450 nm absorbance and subtracted at 540-570 nm. The values were transformed using a four-parameter logistic regression ($R^2$>0.99) to calculate the concentration of FLAMOD in the biological samples.

*Gene expression*

Cultures were washed with PBS (Sigma) and lysed by pipetting RA1 buffer (Macherey Nagel) complemented with 2% tris (2-carboxyethyl) phorphorine (TCEP, Macherey Nagel). Total RNA was then extracted with the Nucleospin RNA kit (Macherey Nagel) and was reverse-transcribed with the High-Capacity cDNA Archive Kit (Applied Biosystems). The cDNA was amplified using TaqMan or SYBR green- based real-time PCR on a Quantstudio 12K PCR system (Applied Biosystems). Relative mRNA levels (2−ΔΔCq) were determined by comparing first the PCR cycle thresholds (Cq) for the gene of interest and the reference genes *Actb* and *B2m* (ΔCq), and then the ΔCq values for FLAMOD-treated vs. vehicle-treated or untreated group (ΔΔCq). All the Taqman assays and primers used in this study are listed in **Supplementary Table 2.**

*RNA extraction, sequencing and analysis*

RNA quality was evaluated using the Fragment Analyzer (TapeStation 4200, Agilent Technologies), and quantified by spectrophotometry (Nanodrop, Thermo Fischer) and fluorimetry (Qubit, Thermo Fisher). The libraries were prepared using the Illumina Stranded mRNA Prep (Illumina; IDT for Illumina RNA UD Indexes Set B, Ligation), and pooled in an equimolar manner prior to sequencing on an Illumina sequencer in 1x75bp. RNA sequencing data (FASTQ files) were mapped and annotated to the human genomen (Ensembl GRCh38.p14, GCA_000001405.29) using the STAR alignment software (STAR 2.7.10a[1]). Genome scaffolds were generated with 74 nucleotide length. Differentially expressed gene and kinetic analysis was performed using the NeNORM v2 method for data normalization[2,3]. Genes (Ensemble Ids) with CDS p-adjusted value 0.05[4] were considered as differentially expressed. Metascape was used to identify statistically enriched Gene Ontology (GO) biological processes[5]. Sequencing data have been deposited in the National Center for Biotechnology Information Gene Expression Omnibus repository under the accession number GSE313945.

*Mass spectrometry proteomic analysis*

Protein samples were fractionated on a 10% acrylamide SDS-PAGE gel. The gel was briefly stained with Coomassie Blue, and one band for apical wash and cell lysate samples, or three bands for basal medium samples were cut and digested by trypsin as described previously[6]. LC-MS/MS was performed using UltiMate 3000 RSLCnano System (Thermo Fisher Scientific, Waltham, MA, USA) as previously described[7,8]. Peptides were fractionated onto a commercial C18 reversed-phase column (75 µm × 250 mm, 2-µm particle, PepMap100 RSLC column, temperature 35 °C, Thermo

Fisher Scientific, Waltham, USA). Trapping was performed during 4 min at 5 µL/min, with solvent A (98% H2O, 2% acetonitrile and 0.1% formic acid). The peptides were eluted using the solvents A (0.1% formic acid in water) and B (0.1% formic acid in acetonitrile) at a flow rate of 300 nL/min. Gradient separation was 3 min at 3% B, 110 min from 3 to 20% B for apical wash and cell lysate samples and 50 min from 3 to 20% B for basal medium samples, followed by 10 min from 20% to 80% B. The eluted peptides from the C18 column were analyzed by Q-Exactive instruments (Thermo Fisher Scientific). The electrospray voltage was 1.9 kV, and the capillary temperature was 275 °C. Full MS scans were acquired in the Orbitrap mass analyzer over *m*/*z* 400–1200 range with a 70,000 (*m*/*z* 200) resolution. The target value was $3\times10^6$. Fifteen most intense peaks with charge state between 2 and 5 were fragmented in the higher-energy collision-activated dissociation cell with normalized collision energy of 27%, and tandem mass spectrum was acquired in the Orbitrap mass analyzer with a resolution of 17,500 at *m*/*z* 200. The target value was $10^5$. The ion selection threshold was $5\times10^4$ counts, and the maximum allowed ion accumulation times were 250 ms for full MS scans and 100 ms for tandem mass spectrum. Dynamic exclusion was set to 30 s. Raw data collected during nanoLC-MS/MS analyses were processed and converted into *.mgf peak list format with Proteome Discoverer Software 1.4 (ThermoFisher Scientific). MS/MS data was interpreted using search engine Mascot (version 2.4.0, Matrix Science, London, UK) installed on a local server. Searches were performed with a tolerance on mass measurement of 10 ppm for precursor and 0.02 Da for-fragment ions, against a composite target-decoy database (80250*2 total entries) built with a *Homo sapiens* SwissProt database (taxonomy ID 9606, January 2019, 20388 entries) fused with a homemade database of chemokines and antimicrobial peptides (593 entries), the sequences of FLAMOD, recombinant trypsin and classical contaminants (119 entries). Cysteine carbamidomethylation, methionine oxidation, protein N-terminal acetylation and cysteine propionamidation were searched as variable modifications. Up to one trypsin missed cleavage were allowed.

The identification results were imported into Proline software (http://proline.profiproteomics.fr) for validation[9]. Peptide spectrum matches taller than 9 residues and ion scores >10 were retained. The false discovery rate was then optimized to be below 1% at the protein level using the Mascot Modified Mudpit score. Spectral counting analyses were performed with Proline 2.0. The analysis was based on the number of spectra corresponding to peptides uniquely mapping to a given protein, relative to the total number of spectra detected in the sample. When indicated, protein-specific peptide counts were normalized to the total peptide spectral count to account for variations in sample input and processing. The data have been deposited to the ProteomeXchange Consortium via the PRIDE repository with the dataset identifier (number pending).

#### SUPPLEMENTARY REFERENCES

1 Dobin A, Davis CA, Schlesinger F, Drenkow J, Zaleski C, Jha S, *et al.* STAR: ultrafast universal RNA-seq aligner. *Bioinformatics* 2013;**29**:15–21. https://doi.org/10.1093/bioinformatics/bts635.

2 Noth S, Brysbaert G, Benecke A. Normalization using weighted negative second order exponential error functions (NeONORM) provides robustness against asymmetries in comparative transcriptome profiles and avoids false calls. *Genomics Proteomics Bioinformatics* 2006;**4**:90–109. https://doi.org/10.1016/S1672-0229(06)60021-1.

3 ARN_dt. *NeONROMv2*. Codeberg.org. n.d. URL: https://codeberg.org/ARN_dt/NeONROMv2 (Accessed 17 December 2025).

4 Tchitchek N, Dzib JFG, Targat B, Noth S, Benecke A, Lesne A. CDS: a fold-change based statistical test for concomitant identification of distinctness and similarity in gene expression analysis. *Genomics Proteomics Bioinformatics* 2012;**10**:127–35. https://doi.org/10.1016/j.gpb.2012.06.002.

5 Zhou Y, Zhou B, Pache L, Chang M, Khodabakhshi AH, Tanaseichuk O, *et al.* Metascape provides a biologist-oriented resource for the analysis of systems-level datasets. *Nat Commun* 2019;**10**:1523. https://doi.org/10.1038/s41467-019-09234-6.

6 Miguet L, Béchade G, Fornecker L, Zink E, Felden C, Gervais C, *et al.* Proteomic analysis of malignant B-cell derived microparticles reveals CD148 as a potentially useful antigenic biomarker for mantle cell lymphoma diagnosis. *J Proteome Res* 2009;**8**:3346–54. https://doi.org/10.1021/pr801102c.

7 Aruçi E, Saliou J-M, Ferveur J-F, Briand L. Proteomic Characterization of Drosophila melanogaster Proboscis. *Biology* 2022;**11**:1687. https://doi.org/10.3390/biology11111687.

8 Mondemé M, Zeroual Y, Soulard D, Hennart B, Beury D, Saliou J-M, *et al.* Amoxicillin treatment of pneumococcal pneumonia impacts bone marrow neutrophil maturation and function. *J Leukoc Biol* 2024;**115**:463–75. https://doi.org/10.1093/jleuko/qiad125.

9 Perez-Riverol Y, Bai J, Bandla C, García-Seisdedos D, Hewapathirana S, Kamatchinathan S, *et al.* The PRIDE database resources in 2022: a hub for mass spectrometry-based proteomics evidences. *Nucleic Acids Res* 2022;**50**:D543–52. https://doi.org/10.1093/nar/gkab1038.

### Supplementary Figures

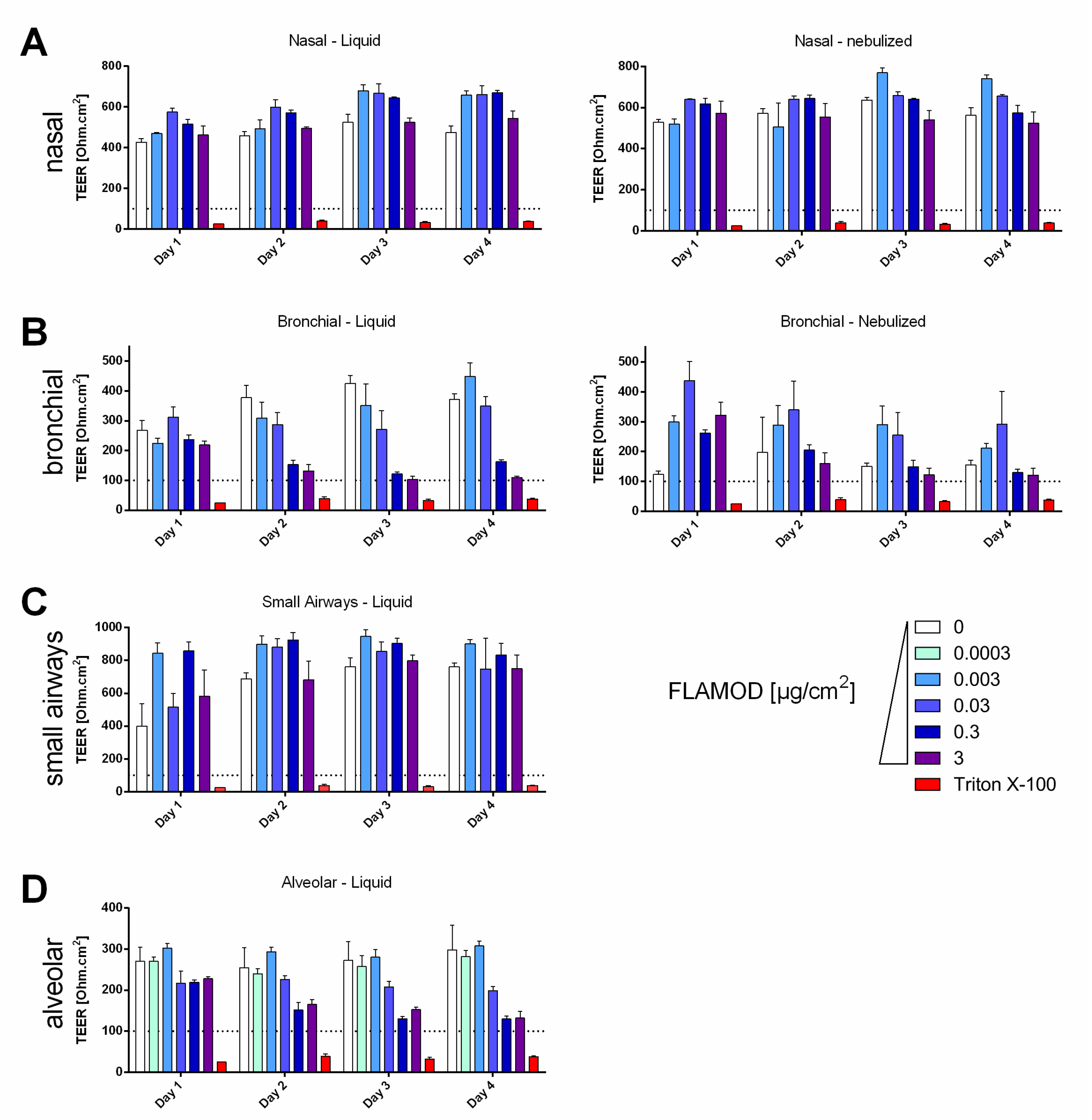

**Supplementary Figure 1. Effect of daily FLAMOD administration on airway epithelium integrity.** Daily exposure to the flagellin FLAMOD started on day 0 and continued for 4 days, as described in Figure 1 (5 exposures). TEER was measured upon FLAMOD pipetting (liquid, left) or nebulization (nebulized, right) in **A)** nasal, **B)** bronchial, **C)** small airways and **D)** alveolar epithelia. Triton X-100 treatment was used as control for total loss of tissue integrity. TEER value < 100 Ω·cm² indicates compromised epithelial barrier function.

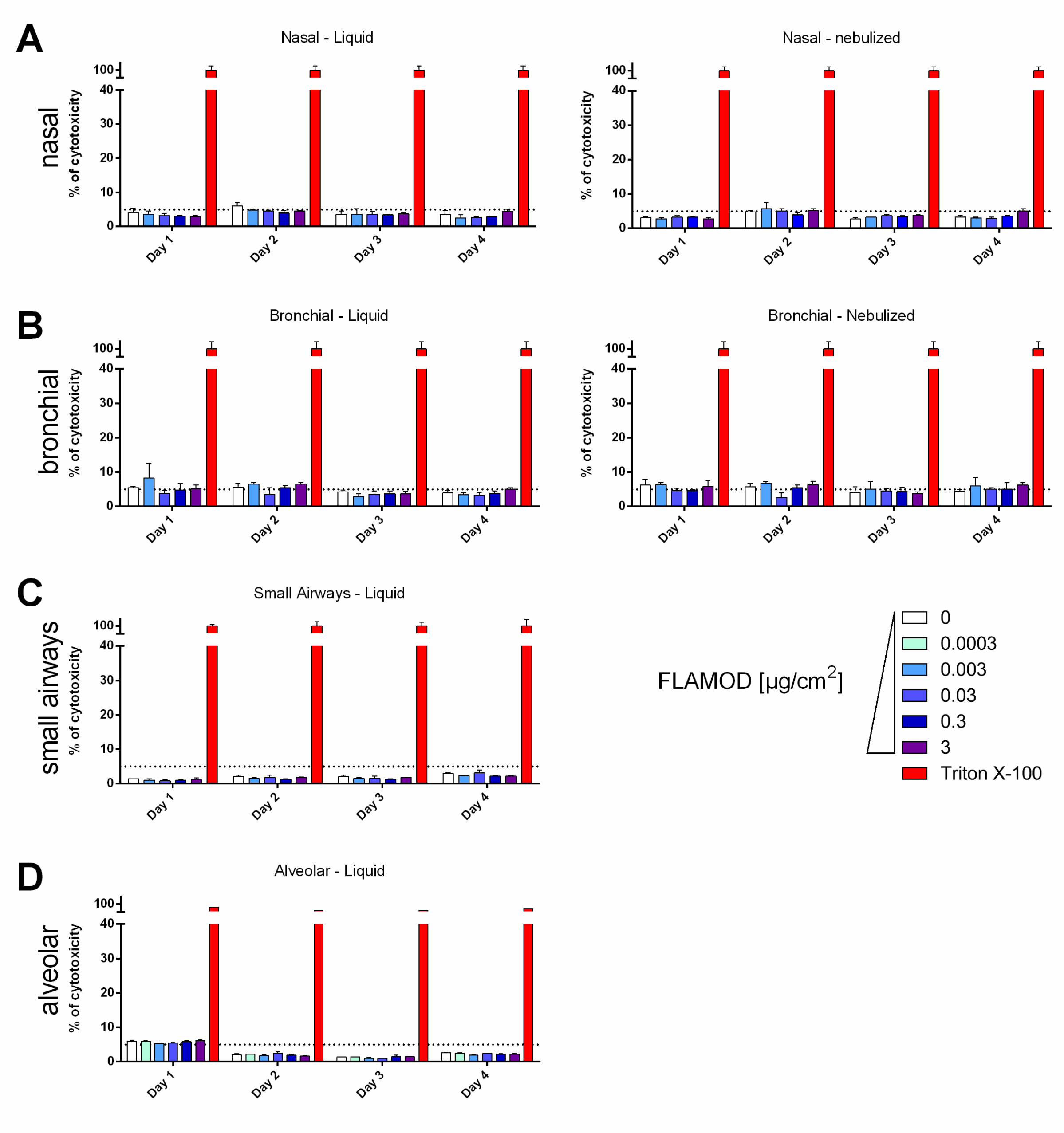

**Supplementary Figure 2. Cytotoxicity of daily FLAMOD administration on airway epithelium.** Daily exposure to the flagellin FLAMOD started on day 0 and continued for 4 days, as described in Figure 1 (5 exposures). LDH release in the basal culture medium was measured upon FLAMOD pipetting (liquid, left) or nebulization (nebulized, right) in **A)** nasal **B)** bronchial, **C)** small airways and **D)** alveolar epithelia. Triton X-100 treatment was used as control for 100% cytotoxicity. A threshold of 5% LDH release that corresponds to the normal level of cell turnover, was used to define the limit of acceptable cytotoxicity N=3 replicates; Data are presented as mean ± SEM.

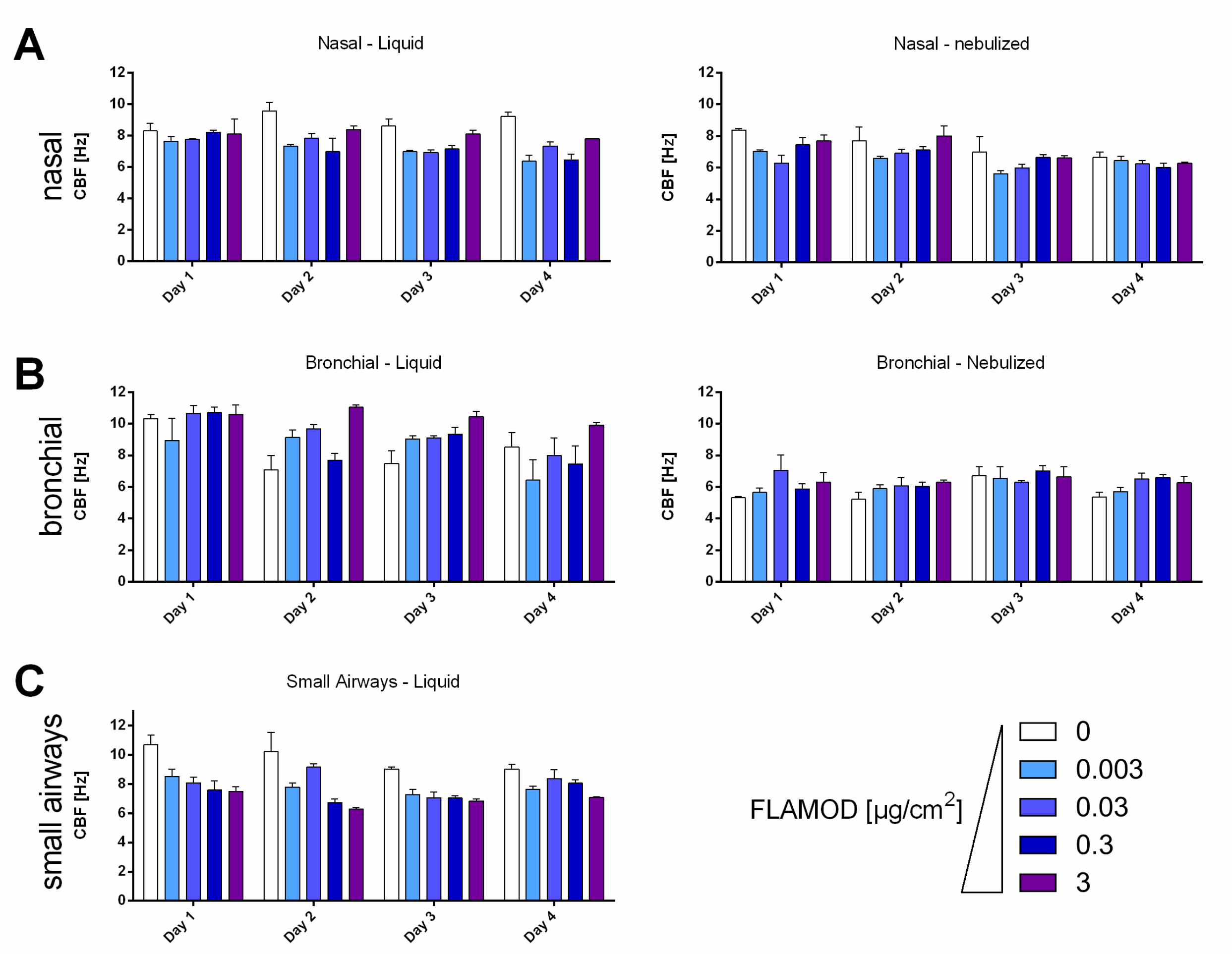

**Supplementary Figure 3. Impact of daily FLAMOD administration on cilia motion.** Daily exposure to the flagellin FLAMOD started on day 0 and continued for 4 days, as described in Figure 1 (5 exposures). Cilia beating frequency (CBF) was measured upon FLAMOD pipetting (liquid, left) or nebulization (nebulized, right) in **A)** nasal, **B)** bronchial, **C)** small airways epithelia**.**

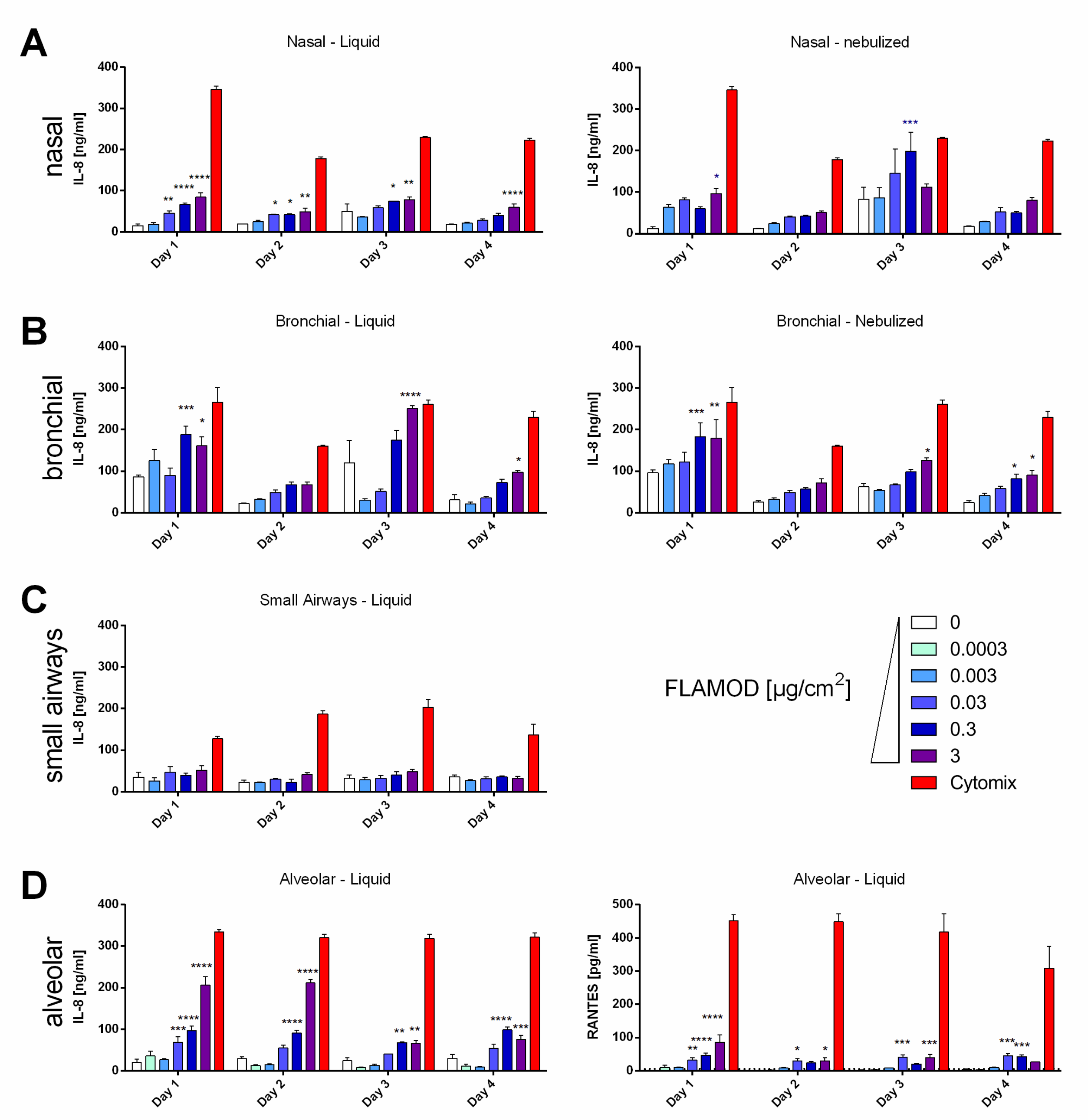

**Supplementary Figure 4. Basal secretion of cytokines/chemokines during daily FLAMOD administration.** Daily exposure to the flagellin FLAMOD started on day 0 and continued for 4 days, as described in Figure 1 (5 exposures). Interleukin-8 (IL-8) secretion in the basal culture medium was measured by ELISA upon FLAMOD pipetting (liquid, left) or nebulization (nebulized, right) in **A)** nasal, **B)** bronchial, **C)** small airways and **D)** alveolar epithelia. Cytomix was used as positive control. RANTES was measured in the basal culture medium of alveolar epithelia (D, right panel). Statistical analysis was performed comparing vehicle control to

FLAMOD-treated tissue using one-way ANOVA with Dunnett's multiple comparison post-test (*$p<0.05$, **$p<0.01$, ***$p<0.001$, ****$p<0.0001$).

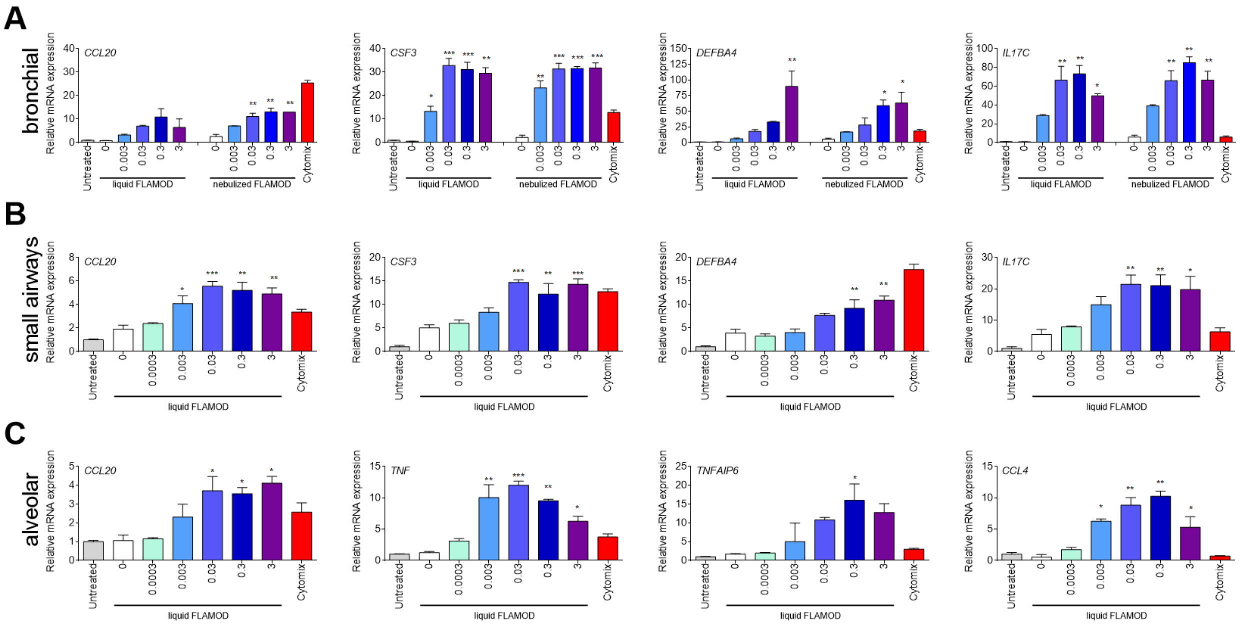

**Supplementary Figure 5. FLAMOD administration by nebulization or liquid droplet efficiently stimulates pro-inflammatory responses in bronchial, small airways and alveolar epithelium**. Daily exposure of respiratory epithelium to FLAMOD (liquid or nebulized on apical interface) was performed as described in Figure 1 (doses in µg/cm$^2$). Epithelia were lysed 2 h after the last exposure for gene expression analysis by RT-qPCR. Cytomix was used to as positive control. Gene expression is presented for **(A)** bronchial, **(B)** small airways and **(C)** alveolar epithelium. Statistical analysis was performed comparing vehicle control to FLAMOD-treated tissue using one-way ANOVA with Dunnett's multiple comparison post-test (*p<0.05, **p<0.01, ***p<0.001, ****p<0.0001).

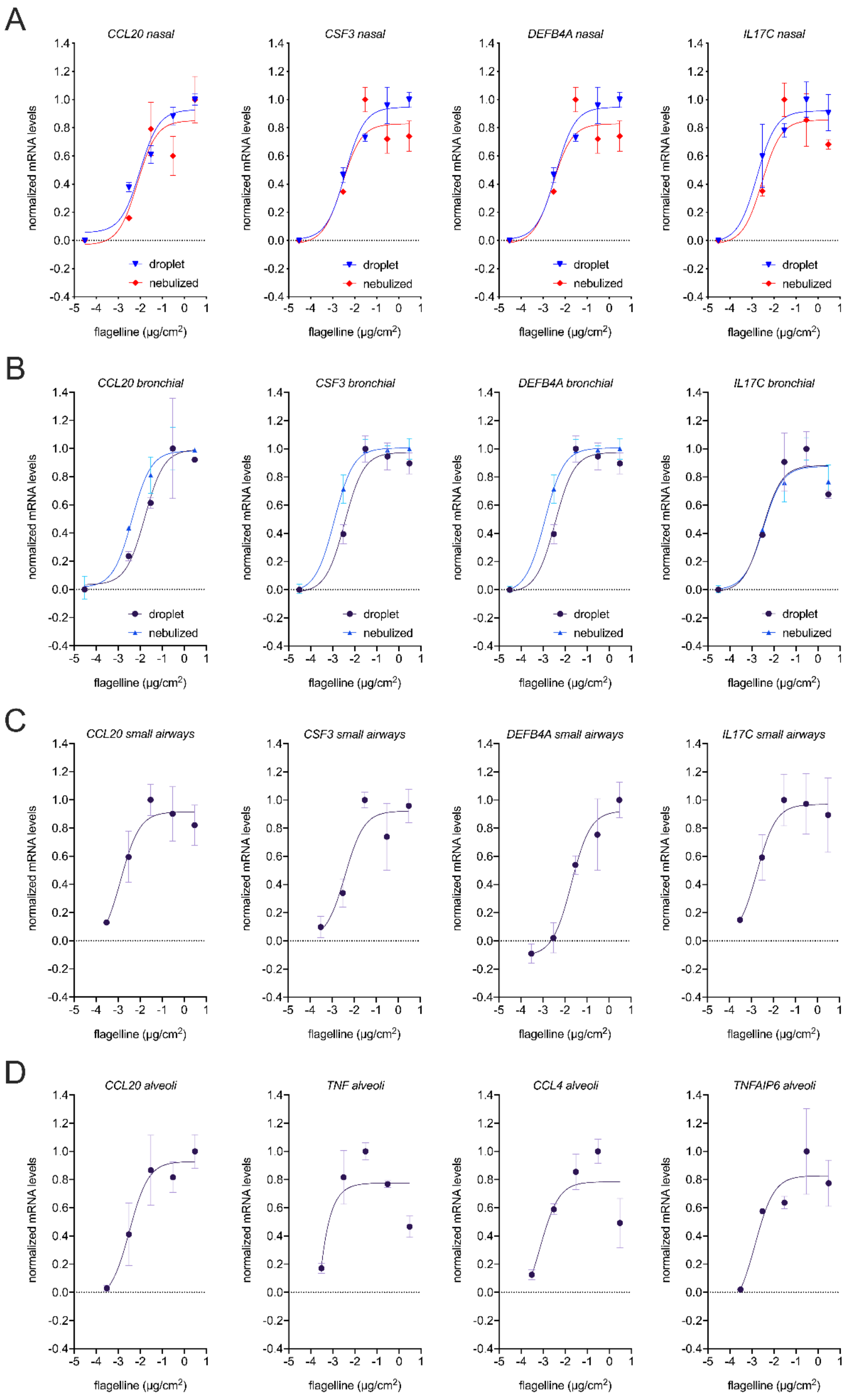
A
CCL20 nasal
CSF3 nasal
DEFB4A nasal
IL17C nasal
droplet
nebulized
B
CCL20 bronchial
CSF3 bronchial
DEFB4A bronchial
IL17C bronchial
droplet
nebulized
C
CCL20 small airways
CSF3 small airways
DEFB4A small airways
IL17C small airways
D
CCL20 alveoli
TNF alveoli
CCL4 alveoli
TNFAIP6 alveoli
normalized mRNA levels
flagelline (µg/cm2)

**Supplementary Figure 6. Analysis of effective dose of FLAMOD in the different human airway epithelium.** Daily exposure of respiratory epithelium to FLAMOD (liquid or nebulized on apical interface) was performed as described in Figure 1 (doses in µg/cm$^2$). Epithelia were lysed 2 h after the last exposure for gene expression analysis by RT-qPCR. Analysis of gene expression after stimulation with various amounts of flagellin. mRNA levels were normalized to house-keeping genes and the condition treated with vehicle was set arbitrarily at the value of 1. A nonlinear fitting of data was performed using a $\log_{10}$(dose of flagellin) vs response curve with three parameters for the indicated genes. The ED50 can be estimated as the dose of agonist that produces 50% of the maximal biological response, i.e., gene expression level. (**A**) Nasal epithelium, (**B**) bronchial epithelium, (**C**) small airway epithelium, and (**D**) alveolar epithelium

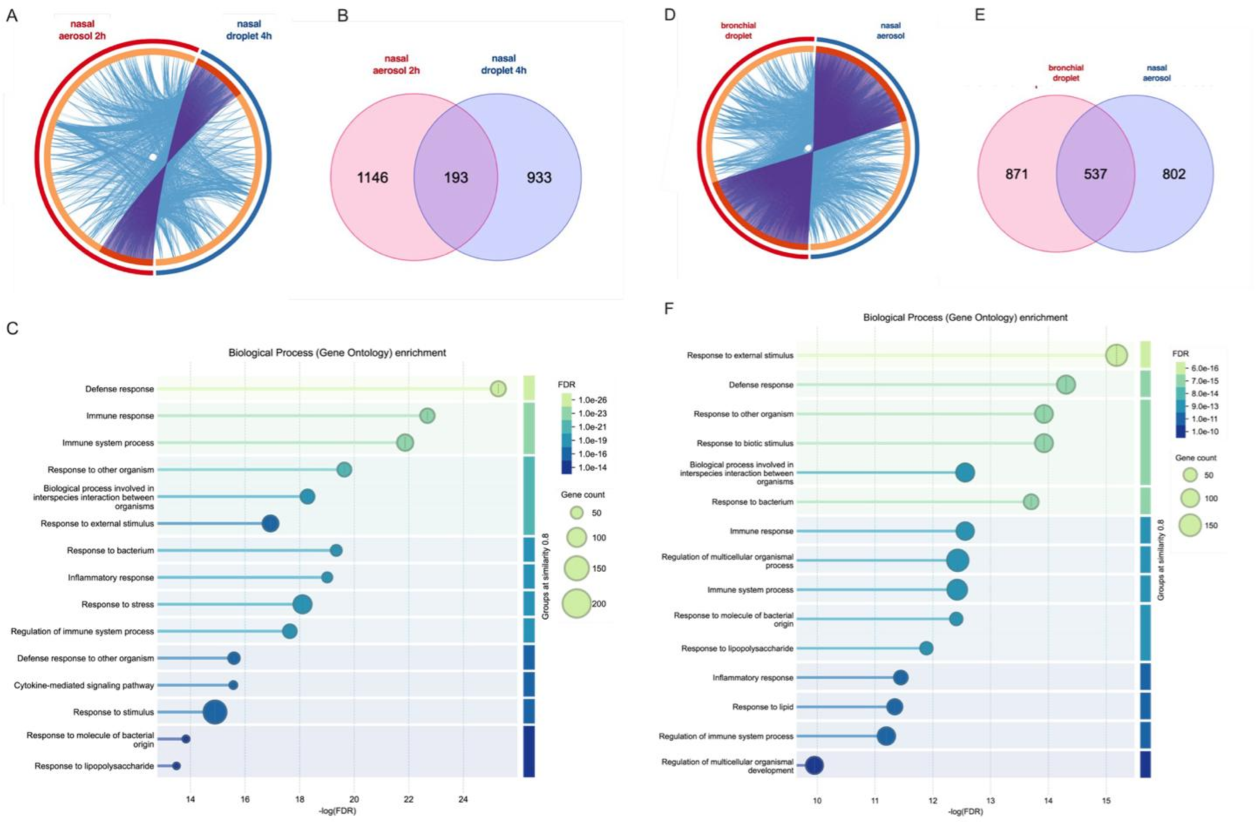

**Supplementary Figure 7**. **Comparative analysis of gene expression patterns across experimental models, delivery routes, and airway compartments.** Total RNA was extracted and processed for RNA-seq from experiments as described in Figure 1A. Samples corresponding to repeated dosing of 0.3 µg/cm$^2$ FLAMOD and vehicle buffer were used to analyze the differentially expressed genes (DEG; listed **in Supplementary File 1**). Metascape (https://metascape.org) was used for gene overlap analysis. (**A, D**) Circos plot depicting gene overlap among indicated conditions. Outer arcs represent each condition, dark orange inner arcs indicate shared genes connected by purple lines, light orange arcs denote unique genes for each condition, and blue lines indicate functional overlap among gene based on shared ontology terms. (**B, E**) Venn diagram illustrating the number of unique and overlapping genes across different conditions. (**C, F**) Graph of enrichment ontology clusters across overlapping genes. (**A-C**) Comparative analysis using a previously generated dataset from nasal epithelium exposed once to droplet for 4 h (GEO accession number GSE304065) and a new dataset from nasal epithelium after 5 repeated aerosol exposures, sampled at 2 h (GEO submission delayed due the lapse in US government funding; data accessible via https://nextcloud.univ-lille.fr/index.php/s/nwRtzwHPGDDHQbk). (D-F) Comparative analysis between bronchial epithelium exposed via droplet and nasal epithelium exposed via aerosol (GEO submission delayed due the lapse in US government funding; data accessible via https://nextcloud.univ-lille.fr/index.php/s/nwRtzwHPGDDHQbk).

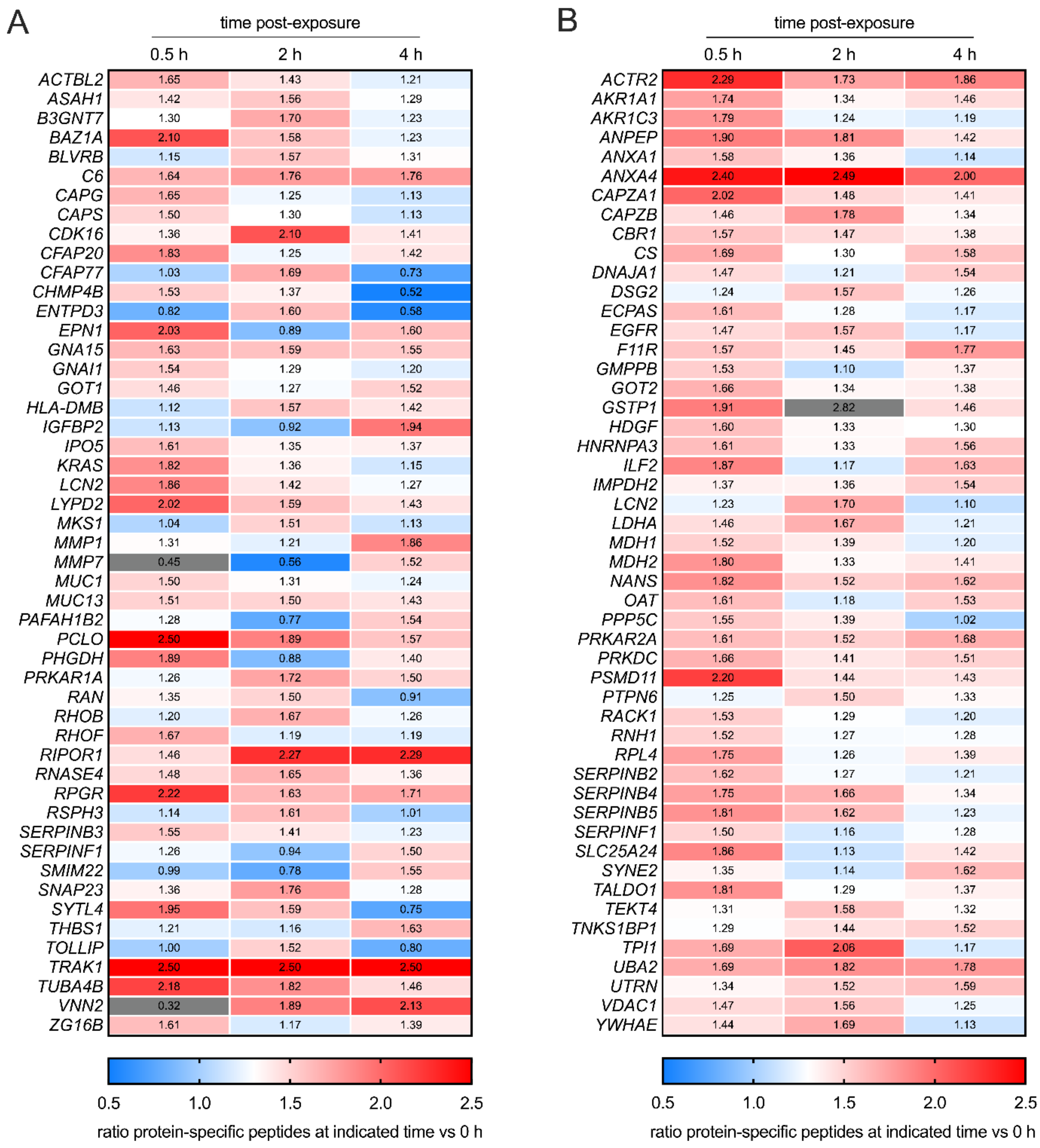

**Supplementary Figure 8. Changes in protein expression associated with FLAMOD treatment in bronchial epithelium.** Primary human bronchial epithelium cultured at the air–liquid interface was exposed for 0 h, 0.5 h, 2 h, and 4 h to FLAMOD applied directly to the apical surface via pipetting as in Figure 3. At the indicated time points post-exposure (n=4 for each time points), apical washes with PBS and lysates of epithelial cells were collected to assess the human protein changes in the apical compartment (**A**), and the whole cell epithelium (**B**), respectively. Proteomic analysis was performed using high-performance liquid chromatography coupled with mass spectrometry to detect protein-specific peptides. Only proteins identified with at least two unique peptides across the 16 samples were included in the analysis. Data were normalized to the total peptide count per sample. Results are presented as heatmaps showing the signal ratio between the mean abundance of specific peptides at each time point and the mean abundance in the control group at 0 h. For clarity, only proteins exhibiting a ratio greater than 1.5 relative to 0 h at any time point are displayed.

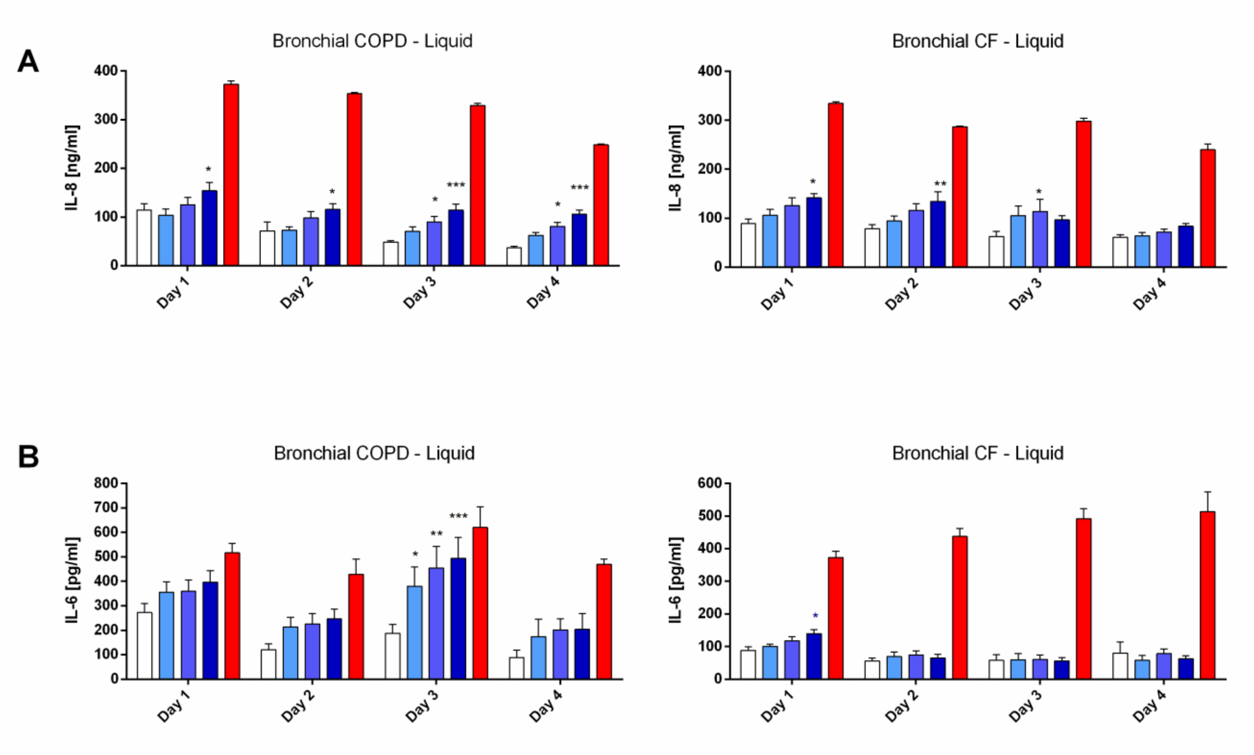

**Supplementary Figure 9. Effect of daily FLAMOD administration in CF and COPD epithelia.** Daily exposure to the flagellin FLAMOD by pipetting (liquid) started on day 0 and continued for 4 days, as described in Figure 1 (5 exposures), using epithelia reconstituted from COPD (left) or CF (right) donors. The effects of daily administration on basal cytokine secretion (**A** interleukin-8 and **B** interleukin-6, positive control is cytomix) was evaluated daily. Statistical analysis was performed comparing vehicle control to FLAMOD-treated tissue using two-way ANOVA with Dunnett's multiple comparison post-test (*p<0.05, **p<0.01, ***p<0.001, ****p<0.0001).

### Supplementary Tables

**Supplementary Table 1. List of chemicals, media, antibodies, kits, equipment, software and deposited data.**

| **Chemicals and media** | | |
|---|---|---|
| **Name** | **Reference** | **Supplier** |
| DPBS (w/o Ca/Mg) | 14190-094 | Gibco, Waltham, MA, USA |
| PBS (with Ca/Mg) | D8662 | Sigma, Burlington, MA, USA |
| Disodium hydrogen phosphate dihydrate | 1.06576 | Merck, Darmstadt, Germany |
| Sodium dihydrogen phosphate dihydrate | 1.37018 | Merck, Darmstadt, Germany |
| Polysorbate 80 | 8.17061 | Merck, Darmstadt, Germany |
| NaCl | 1.37017 | Merck, Darmstadt, Germany |
| NaCl 0.9% | 100 0 266 | Bichsel, Interlaken, Switzerland |
| $H_2O$ (sterile) | B230531 | Versylene Fresenius, Bad Homburg vor der Höhe, Germany |
| $CaCl_2$ | 141221 | ITW Reagent, Castellar del Vallès, Spain |
| TNF | 570104 | Biolegend, San Diego, CA, USA |
| LPS | L9143 | Sigma, Burlington, MA, USA |
| FCS | 2-01F16-I | Bioconcept, Allschwil, Switzerland |
| HEPES | 15030080 | Gibco, Waltham, MA, USA |
| $H_2SO_4$ | 07208 | Sigma, Burlington, MA, USA |
| Tween 20 | A4974 | ITW Reagent, Castellar del Vallès, Spain |
| Triton-X100 | 93418 | Fluka, Buchs, Switzerland |
| RA1 buffer | 11912412 | Macherey Nagel, Düren, Germany |
| TCEP | 12703401 | Macherey Nagel, Düren, Germany |
| TMB Substrate | UP664781 | Interchim, Montluçon, France |
| ELISA/ELISPOT diluent 5X | 00-4202-56 | Invitrogen, Waltham, MA, USA |
| Avidin-HRP | 18-4100-51 | eBioscience, San Diego, CA, USA |
| Antibodies | | |
| **Name** | **Reference** | **Supplier** |
| Capture anti-FLAMOD (mAb) | 9H10-R2-2E8<br>mouse IgG1 | Biotem, France |
| Detection anti-FLAMOD (mAb) | 4C1H7<br>mouse IgG1 | Biotem, France |

| Kits | | |
|---|---|---|
| **Name** | **Reference** | **Supplier** |
| HEK-Dual™ hTLR5 cells assay | hkd-htlr5ni | Invivogen, San Diego, CA, USA |
| Cytotoxicity LDH Assay Kit-WST | CK12-20 | Dojindo, Mashiki, Japan |
| Pyrochrome® | NC1241122 | Associates of Cape Cod Inc., East Falmouth, MA, USA |
| Human IL-8 ELISA Set | 555244 | BD Biosciences, Franklin Lakes, NJ, USA |
| Human CCL5/RANTES DuoSet ELISA | DY278 | R&D Systems, Minneapolis, MN, USA |
| Nucleospin RNA kit | 740955 | Macherey Nagel, Düren, Germany |
| High-Capacity cDNA Archive Kit | 4368814 | Applied Biosystems, Foster City, CA, USA |
| Takyon Low Rox SYBR 2X MasterMix | UF-LSMT-B0710 | Eurogentec, Seraing, Belgium |
| Takyon Low ROX Probe 2X MasterMix | UF-LPMT-B0705 | Eurogentec, Seraing, Belgium |
| TaqMan array 96-well fast plate | 4413261 | Thermo Fisher Scientific, Waltham, MA, USA |
| Illumina® RNA UD Indexes Set D, Ligation | 20091661 | Illumina, San Diego, CA, USA |
| Equipment | | |
| **Name** | **Function** | **supplier** |
| Aerogen Solo (AG-AS3200) with A-VMN™ mesh | Mesh Nebulizer | Aerogen, Paris, France |
| Aerogen USB controller (AGUC1000NE) | USB controller | Aerogen, Paris, France |
| Maxisorp 96 well plate | ELISA plate (442404) | Nalgen Nunc International, Roskilde, Denmark |
| Multiskan FC | ELISA plate reader | Thermo Fisher Scientific, Waltham, MA, USA |
| UltiMate 3000 RSLCnano | Liquid chromatography | Thermo Fisher Scientific, Waltham, MA, USA |
| Orbitrap Mass Spectrometer | Mass analyzer | Thermo Fisher Scientific, Waltham, MA, USA |
| VITROCELL® Cloud alpha 12 | Aerosolization | Vitrocell, Waldkirch, Germany |
| EVOMX volt-ohm-meter | Resistance measurement | World Precision Intrument, Sarasota, FL, USA |
| Victor Nivo | Absorbance plate reader | Perkin Elmer, Waltham, MA, USA |
| Mako G030B | High-speed camera | Allied Vision Technologies, Cupertino, CA, USA |

| | | |
|---|---|---|
| Quantstudio 12K PCR system | Thermal cycler | Applied Biosystems, Foster City, CA, USA |
| 2100 Bioanalyzer | Spectrophotometer | Agilent Technologies, Santa Clara, CA, USA |
| NovaSeq 2000 system | High throughput sequencer | Illumina, San Diego, CA, USA |
| TapeStation 4200 | Fragment Analyzer | Agilent Technologies, Santa Clara, CA, USA |
| Qubit | Fluorometer | Thermo Fisher Scientific, Waltham, MA, USA |
| Software | | |
| Cilia-X | CBF calculation | Epithelix Sàrl, Geneva, Switzerland |
| Prism 6 and 8.4.3 GraphPad | Graph statistical analysis | GraphPad Software, La Jolla, CA, USA |
| STAR alignment 2.7.10a | RNA-seq aligner | Dobin *et al.* https://pmc.ncbi.nlm.nih.gov/articles/PMC3530905/ |
| DESeq2 | Gene expression | Bioconductor |
| NeONORM | Gene expression | Noth *et al.* https://github.com/systemsepigenomics/neonorm |
| Deposited data | | |
| RNA-seq data | This paper - Gene Expression Omnibus | Due to the lapse of US funding, GEO submission will be done later. Data are however available on https://nextcloud.univ-lille.fr/index.php/s/nwRtzwHPGDDHQbk |

**Supplementary Table 2. Epithelium characteristics.**

| Epithelium characteristics | | | |
|---|---|---|---|
| **Figure** | **Epithelium** | **Donor** | **Pathology*** |
| Figure 1 | Nasal | Pool | No |
| Figure 2 | Nasal | Pool | No |
| Figure 3 | Bronchial | Unique | No |
| Figures 4 and 5 | Bronchial | Unique | COPD or CF |
| Supplementary Figure 1 | Nasal<br>Bronchial<br>Small airways<br>Alveolar | Pool<br>Unique<br>Unique<br>Unique | No<br>No<br>No<br>No |
| Supplementary Figure 2 | Nasal<br>Bronchial<br>Small airways<br>Alveolar | Pool<br>Unique<br>Unique<br>Unique | No<br>No<br>No<br>No |
| Supplementary Figure 3 | Nasal<br>Bronchial<br>Small airways | Pool<br>Unique<br>Unique | No<br>No<br>No |
| Supplementary Figure 4 | Nasal<br>Bronchial<br>Small airways<br>Alveolar | Pool<br>Unique<br>Unique<br>Unique | No<br>No<br>No<br>No |
| Supplementary Figure 5 | Bronchial<br>Small airways<br>Alveolar | Unique<br>Unique<br>Unique | No<br>No<br>No |
| Supplementary Figure 6 | Nasal<br>Bronchial<br>Small airways<br>Alveolar | Pool<br>Unique<br>Unique<br>Unique | No<br>No<br>No<br>No |
| Supplementary Figure 7 | Nasal<br>Bronchial | Pool<br>Unique | No<br>No |
| Supplementary Figure 8 | Bronchial | Unique | No |
| Supplementary Figure 9 | Bronchial | Unique | COPD or CF |

* No stand for healthy donor

**Supplementary Table 3. List of assays and primers.**

| Gene | full name | TaqMan assay number | Sybr green assay forward primer | Sybr green assay reverse primer |
|---|---|---|---|---|
| ***ACTB*** | actin beta | Hs03023943_g1 | ATTGGCAATGAGCGGTTC | CGTGGATGCCACAGGACT |
| ***B2M*** | beta-2-microglobulin | Hs00187842_m1 | TTCTGGCCTGGAGGCTATC | TCAGGAAATTTGACTTTCCATTC |
| ***CCL20*** | C-C motif chemokine ligand 20 | Hs00355476_m1 | CCAAGAGTTTGCTCCTGGCT | TGCTTGCTGCTTCTGATTCG |
| ***CCL4*** | C-C motif chemokine ligand 4 | n/a | CTTCCTCGCAACTTTGTGGT | CAGCACAGACTTGCTTGCTT |
| ***CSF3*** | colony stimulating factor 3 | Hs00357085_g1 | GTGCTGCTCGGACACTCTCT | GAAAAGGCCGCTATGGAGTT |
| ***CXCL8*** | C-X-C motif chemokine ligand 8 (interleukin 8) | n/a | CACCGGAAGGAACCATCTCA | GGAAGGCTGCCAGAGAGC |
| ***CYP3A5*** | cytochrome P450 family 3 subfamily A member 5 | Hs00241417_m1 | n/a | n/a |
| ***CYP3A7*** | cytochrome P450 family 3 subfamily A member 7 | Hs00426361_m1 | n/a | n/a |
| ***DEFB4A*** | defensin beta 4A | Hs00175474_m1 | TCAGCCATGAGGGTCTTGTA | AGGATCGCCTATACCACCAA |
| ***EBI3*** | Epstein-Barr virus induced 3 | Hs01057148_m1 | n/a | n/a |
| ***IDO1*** | indoleamine 2,3-dioxygenase 1 | Hs00984148_m1 | CAAAGCAGCGTCTTTCAGTG | AGGAACTGAGCAGCATGTCC |
| ***IL17C*** | interleukin 17C | Hs00171163_m1 | CCCTCAGCTACGACCCAGT | CTTCTGTGGATAGCGGTCCT |
| ***IL1B*** | interleukin 1 beta | n/a | TACCTGTCCTGCGTGTTGAA | TCTTTGGGTAATTTTTGGGATCT |
| ***LRRC55*** | leucine rich repeat containing 55 | Hs01590358_m1 | CTGCAGATCGGTGGCAAT | CAGCCAGCTGAGAATCTGC |
| ***PGLYRP2*** | peptidoglycan recognition protein 2 | Hs00994650_m1 | n/a | n/a |
| ***TNF*** | tumor necrosis factor | n/a | CCCAGGCAGTCAGATCATCTTC | GGTTTGCTACAACATGGGCTACA |
| ***TNFAIP6*** | tumor necrosis factor alpha induced protein 6 | n/a | GGCCATCTCGCAACTTACA | GCAGCACAGACATGAAATCC |
| ***TNFRSF6B*** | TNF receptor superfamily member 6b | Hs00187070_m1 | n/a | n/a |
| ***TNIP3*** | TNFAIP3 interacting protein 3 | Hs00375573_m1 | n/a | n/a |
| ***UBASH3A*** | ubiquitin associated and SH3 domain containing A | Hs00955168_m1 | n/a | n/a |
